# Multi-hallmark long noncoding RNA maps reveal non-small cell lung cancer vulnerabilities

**DOI:** 10.1101/2021.10.19.464956

**Authors:** Roberta Esposito, Taisia Polidori, Dominik F. Meise, Carlos Pulido-Quetglas, Panagiotis Chouvardas, Stefan Forster, Paulina Schaerer, Andrea Kobel, Juliette Schlatter, Michaela Roemmele, Emily S. Westemeier, Lina Zhu, Andrés Lanzós, Hugo A. Guillen-Ramirez, Giulia Basile, Irene Carrozzo, Adrienne Vancura, Sebastian Ullrich, Alvaro Andrades, Dylan Harvey, Pedro P. Medina, Patrick C. Ma, Simon Haefliger, Xin Wang, Ivan Martinez, Adrian Ochsenbein, Carsten Riether, Rory Johnson

## Abstract

Long noncoding RNAs (lncRNAs) are widely dysregulated in cancer, yet their functional roles in cellular disease hallmarks remain unclear. Here we employ pooled CRISPR deletion to perturb all 831 lncRNAs in KRAS-mutant non-small cell lung cancer (NSCLC), and measure their contribution to proliferation, chemoresistance and migration across two cell backgrounds. Integrative analysis of this data outperforms conventional “dropout” screens in identifying cancer genes, while prioritising disease-relevant lncRNAs with pleiotropic and background-independent roles. Altogether 60 high-confidence oncogenic lncRNAs are active in NSCLC, the majority identified here for the first time, and which tend to be amplified and overexpressed in tumours. A follow-up antisense oligonucleotide (ASO) screen shortlisted two candidates, Cancer Hallmarks in Lung LncRNA (CHiLL 1&2), whose knockdown consistently suppressed cancer hallmarks in a variety of 2D and 3D tumour models. Molecular phenotyping reveals that CHiLL 1&2 control cellular-level phenotypes via distinct transcriptional networks converging on common oncogenic pathways. In summary, this work reveals a multi-dimensional functional lncRNA landscape underlying NSCLC that contains potential therapeutic vulnerabilities.

## Introduction

Non-small cell lung cancer (NSCLC) is the leading cause of cancer deaths worldwide (1), and available therapies face a combination of challenges in undruggable mutations, toxicity and therapy resistance (2–4). The most common subtype, carrying activating *KRAS* mutations (KRAS^+^), is routinely treated with cytotoxic platinum chemotherapy, and newly-approved targeted therapies only extend life by few months (5,6).

A fertile source for new therapeutic targets is long noncoding RNAs (lncRNAs), with a population likely to exceed 100,000, of which >98% remain uncharacterised (7–10). Hundreds of lncRNAs have been implicated in disease hallmarks across cancer types via a variety of mechanisms (11–14). Examples such as SAMMSON (melanoma) and lncGRS-1 (glioma) have attracted attention as drug targets, thanks to tumour cells’ potent and specific sensitivity to their inhibition via antisense oligonucleotide (ASO) therapies (15,16). Nonetheless, the extent and nature of lncRNAs promoting the interlocking pathological hallmarks in a given tumour remains unclear.

Efforts to identify lncRNA therapeutic targets have accelerated with the advent of CRISPR-Cas genome-editing, which can be used to silence gene expression via targeted genomic deletions or transcriptional inhibition, and is readily scaled transcriptome-wide via pooling (17). CRISPR screens have revealed scores of lncRNAs promoting disease hallmarks of cell proliferation, pathway activation and therapy resistance (18–20).

A critical challenge in drug discovery is the poor validation rate of preclinical targets discovered *in vitro* (21). Highly-focussed single background / single hallmark screen designs, including those above, are vulnerable to discovering lncRNAs promoting cell-line specific phenotypes that do not generalise to the disease in question (15). Supporting this, recent CRISPR-inhibition (CRISPRi) screens demonstrated highly specific effects for lncRNAs in cell lines from six distinct cancer types (18). However, the critical question of whether this also affects cell lines from the same cancer type has not been addressed. Thus, to maximise the utility of discovered hits, an ideal screen should prioritise lncRNA targets that are both pleiotropic (impact multiple disease hallmarks) and background-independent (effective regardless of cell model).

Here, we comprehensively map the functional lncRNA landscape of *KRAS^+^* NSCLC. We perform 10 disease-specific CRISPR screens for lncRNAs promoting three cancer hallmarks in two cell models. We reconstruct the functional lncRNA landscape of NSCLC, revealing a catalogue of therapeutic vulnerabilities. These lncRNAs connect to cancer hallmarks via complex transcriptional networks, and can be targeted by potent, low toxicity and on-target ASOs, representing promising future therapeutics (22).

## Results

### A versatile CRISPR screening pipeline for long non-coding RNAs in NSCLC

To identify lncRNAs promoting KRAS^+^ NSCLC, we adapted the DECKO (dual excision CRISPR knock-out) CRISPR-deletion (CRISPR-del) system to high-throughput pooled format (17,23) (Figure 1a). This approach achieves loss-of-function perturbations by deleting target genes’ transcription start site (TSS) via paired guide RNAs (pgRNAs), and effectively inhibits gene expression (24–27).

**Figure 1.**
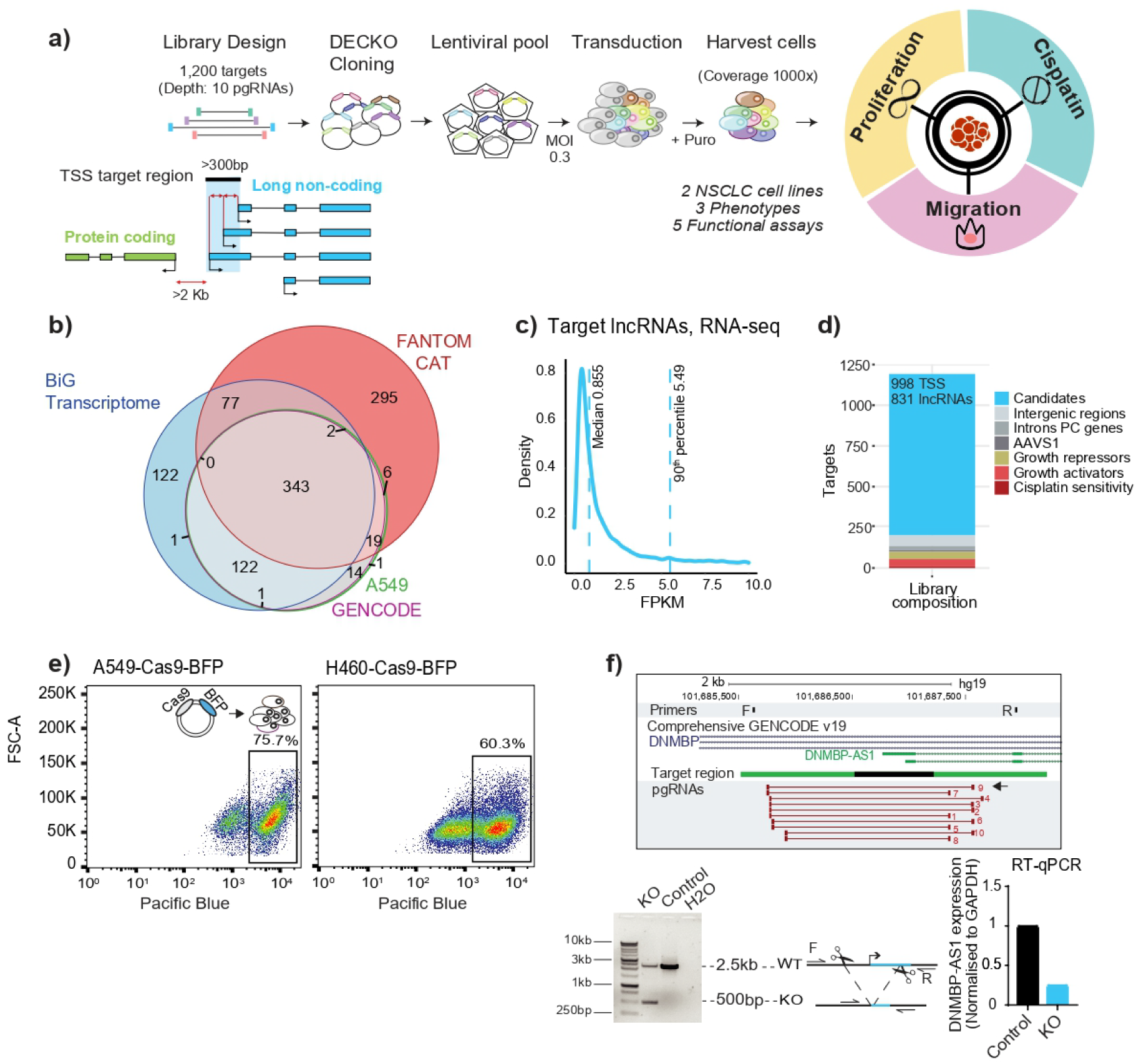
Multi-hallmark CRISPR discovery of lncRNAs promoting non-small cell lung cancer. **a)** CRISPR-deletion pooled screening strategy for lncRNAs promoting NSCLC hallmarks. **b**) Gene annotations used for candidates’ selection. Numbers indicate lncRNA gene loci. **c)** Expression of targeted lncRNAs in A549 cells. **d)** Library composition, in terms of targeted regions. Note that some lncRNA loci are represented by >1 targeted transcription start site (TSS). **e)** Fluorescence activated cell sorting (FACS) was used to sort stable Cas9 expressing cells based on expression of a Blue Fluorescent Protein (BFP) marker. Boxes indicated the sorted cell populations used in screens. **f**) One member of the screening library, DNMBP-AS1, was targeted by CRISPR-deletion in A549 cells. The gene locus is shown in the upper panel, including genotyping PCR primers (F, R), transcription start site (TSS) target region (black), and library paired guide RNAs (pgRNAs, red bars). The pgRNA used here is indicated by the arrow. Below left: PCR using indicated primers with template genomic DNA (gDNA) from cells transfected with non-targeting pgRNA (Control) or DNMBP-AS1 TSS pgRNA (KO). The expected lengths for wild-type and deletion amplicons are indicated. Below right: Quantitative reverse transcriptase PCR (RT-PCR) measurement of DNMBP-AS1 RNA.

We developed a screening library to comprehensively interrogate the NSCLC lncRNA transcriptome. We integrated and filtered published (10,28,29) and in-house annotations (30,31) (Figure 1b), for a final target set of 831 lncRNAs, corresponding to 998 high-confidence TSSs, henceforth named “Candidate_1” and so on (Figure S1a). (Figure 1c and 1d; see Methods). To these we added pgRNAs targeting neutral control loci (not expected to influence cell phenotype) and positive control protein-coding genes (PCGs) (with known roles in cell proliferation and cisplatin resistance) (Figure 1d).

These targets form the basis for ‘libDECKO-NSCLC1’, a CRISPR-deletion library with a depth of 10 unique pgRNAs per target, comprising altogether 12,000 pgRNAs (Figure 1d and File S1). After cloning into the DECKO backbone (Figure S1b), sequencing revealed high quality in terms of sequence identity (60.8% perfect match across both spacers) and coverage (90^th^ to 10^th^ percentile count ratios: 4.6-fold) (32) (Figure S1c).

To identify hits of general relevance to NSCLC, we performed parallel experiments in two widely-used KRAS^+^ NSCLC models, A549 and H460(33,34). Non-clonal cell lines were generated that stably express high-levels of Cas9 protein (35), as evidenced by blue fluorescent protein (BFP) (Figure 1e). Targeting known NSCLC-promoting lncRNA DNMBP-AS1 (Candidate_331) (36), resulted in deletion of its promoter region and loss of expression (Figure 1f), supporting the effectiveness of the CRISPR-del strategy.

### Multi-phenotype mapping of NSCLC lncRNAs

Cancers thrive via a variety of phenotypic “hallmarks” (37). Previous CRISPR screens have been limited to a single hallmark, either proliferation or drug resistance (15,18,38–40), and usually focussed on a negative “drop-out” format, where pgRNAs for genes of interest are depleted.

For more comprehensive and biomedically-relevant vista of NSCLC lncRNAs, we adapted pooled screening to read out distinct hallmarks of proliferation, chemo-resistance and invasion (Figures 2a-c). To boost sensitivity, we implemented complementary “positive” screens, where pgRNAs of interest are enriched. Thus, to identify lncRNAs promoting cell fitness and proliferation, we combined (i) a classical drop-out, where targets’ pgRNAs become depleted, and (ii) a positive screen using CFSE (Carboxyfluorescein succinimidyl ester) dye to identify growth-promoting lncRNAs by their pgRNAs enrichment in slow-growing cells (Figure 2a) (41).

**Figure 2.**
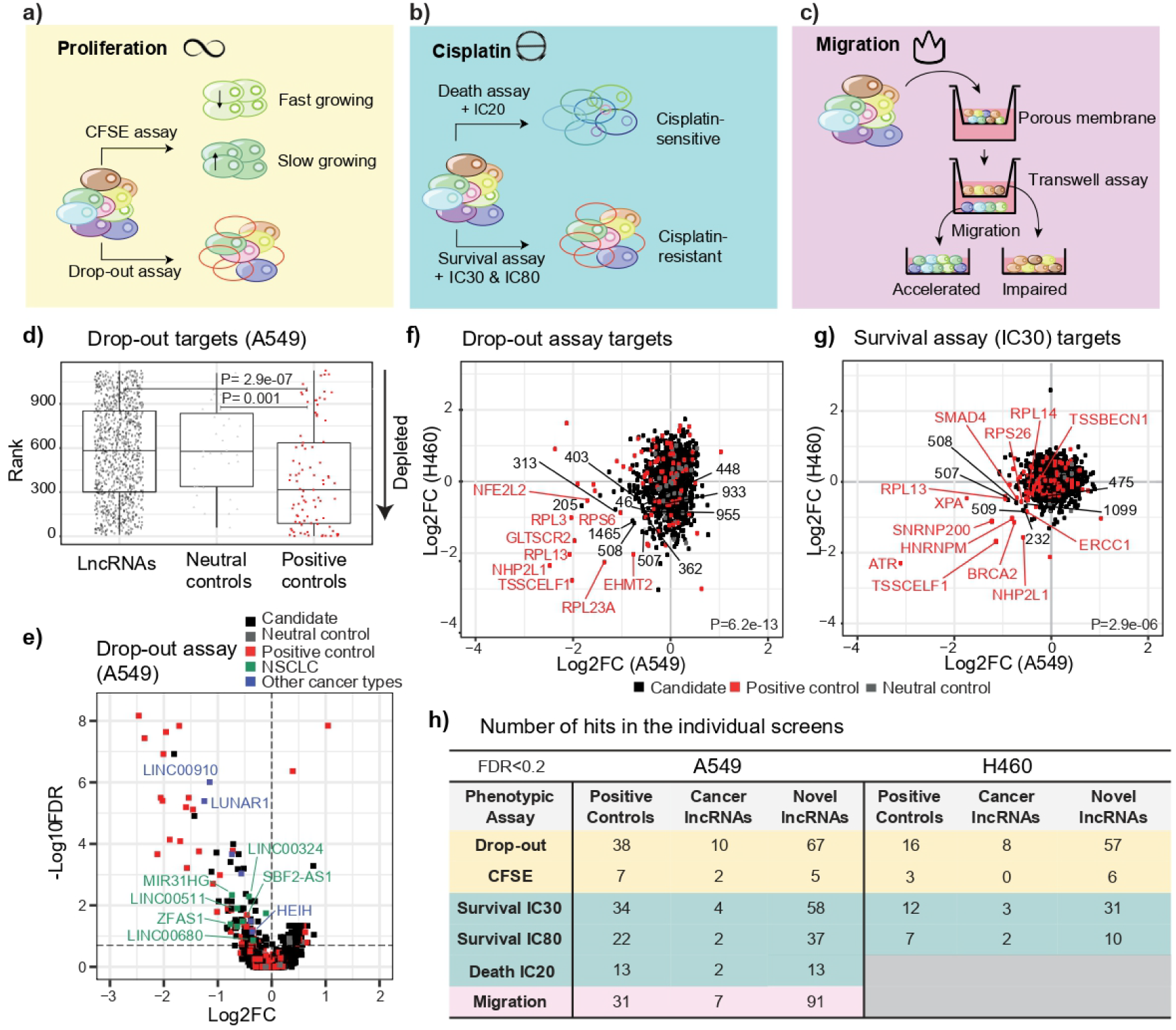
Adapting CRISPR screens to cancer hallmarks. **a)** Proliferation: The strategy employs complementary negative (drop-out) (growth-promoting lncRNAs’ pgRNAs are depleted) and positive (CFSE dye) (growth-promoting lncRNAs’ pgRNAs are enriched) formats. **b)** Cisplatin sensitivity: Another complementary strategy is employed. In the negative (drop-out) “survival” screen, cells are exposed to high cisplatin doses (IC30, IC80). Resistance-promoting lncRNAs’ pgRNAs will be depleted in surviving cells. In the positive “death” screen, cells that die in response to low cisplatin concentration (IC20) are collected, and enriched pgRNAs identify resistance-promoting lncRNAs. **c)** Migration: Cells that are capable / incapable of migrating through a porous membrane over a given time period are separately collected. Migration-promoting lncRNAs are identified via their pgRNAs’ enrichment in migration-impaired cells. **d)** LncRNA candidates, neutral and positive controls, ranked by P-value in A549 drop-out screen (statistical significance estimate using Wilcoxon test). **e)** A549 drop-out screen. Horizontal line indicates cutoff for hits at FDR<0.2. Previously published lncRNAs in NSCLC and other cancers are labelled in green and blue, respectively. **f**) Comparison of A549 and H460 drop-out screens (statistical significance estimated using Pearson correlation). **g**) Comparison of A549 and H460 cisplatin survival screen (Pearson correlation). **h**) Numbers of screen hits at FDR<0.2

As expected, pgRNAs for positive-control genes were significantly depleted in drop-out screens, while neutral controls were not (Figure 2d). To gauge the LOF efficiency of promoter deletion, pgRNAs targeting positive-control protein-coding genes had been split between two distinct modalities: (1) conventional open reading frame (ORF) mutation, expected to yield maximal LOF; and (2) promoter-deletion, similar to lncRNAs. Promoter-deletion pgRNAs displayed a detectable but lower phenotypic impact, indicating that a CRISPR-del screens for lncRNAs face intrinsically lower sensitivity compared to ORF-targeting screens for PCGs (Figure S2a).

Using biologically-replicated drop-out screens, 77 lncRNAs were identified as necessary for proliferation of A549 cells. These include known NSCLC lncRNAs, such as *LINC00324* (42), *ZFAS1* (43), *MIR31HG* (44), *SBF2-AS1* (45), *LINC00680* (46) and *LINC00511* (47), that was also found in H460. In addition, lncRNAs identified in other cancer types include *LUNAR1* (48), *HEIH* (49) and *LINC00910* (50) (Figure 2e).

The factors influencing pgRNA deletion efficiency are poorly understood. Using growth phenotype as a proxy for deletion efficiency, we observed expected correlation between observed and bioinformatically-predicted sgRNA efficiency (RuleSet2 algorithm) (Figure S1d). On the other hand, we found no relationship with pgRNA orientation (Figure S1e), and a weak tendency for larger deletions to produce stronger phenotypes, possibly due to greater impact on lncRNA expression (Figure S1f).

Next, we compared equivalent drop-out screens in the two NSCLC backgrounds. There was a significant concordance amongst identified targets, driven mainly by positive controls (Figure 2f). To strengthen these data, we performed complementary CFSE screens in the same cells. As expected, these positive screens displayed anti-correlation with drop-out results in H460 cells, although not in A549 cells, possibly for technical reasons (Figure S2b).

Patients with KRAS^+^ tumours are usually treated with cytotoxic platinum-based chemotherapeutics, but tumours frequently evolve resistance (51). To identify lncRNAs promoting chemoresistance, we again employed complementary screens (Figure 2B) at carefully chosen cisplatin concentrations (Figures S2d and S2e). As before, correlated results were observed across cell backgrounds (Figure 2g), and PCGs with known roles in cisplatin resistance were correctly identified (red points in Figure 2g). As expected, complementary survival (negative) and death (positive) screens were anti-correlated for high cisplatin dose (IC80) (Figure S2c).

Migration is a key hallmark underlying invasion and metastasis of tumour cells. By isolating cells with rapid or slow migration through a porous membrane for 48h (Figures 2c and S2f), we screened for migration-promoting lncRNAs. This yielded 98 lncRNAs, of which seven were already associated with migration and invasion in numerous cancer types (11), including *NORAD*, *DANCR* and *SNHG29* (52–55).

These data, summarized in Figure 2h and File S2, represent a resource of functional lncRNAs in NSCLC hallmarks.

### Screen hits can be validated and function via RNA products

We next tested the reliability of these results by selecting two lncRNA TSSs for further validation, based on their top ranking and consistency between the two cell lines: Candidate_205, identified as top hit in drop-out screens (A549: Log2FC=-1.81, FDR=1.2e-07) and Candidate_509 in both proliferation and cisplatin (A549: Log2FC=-1.15, FDR=9.78e-07).

Candidate_205 overlaps the TSS of bidirectional antisense GENCODE-annotated genes, *LINC00115* and *RP11-206L10* (Figure 3a). Candidate_509 targets a TSS shared by several BIGTranscriptome lncRNAs (Figure 3a). Supporting the importance of this locus, it contains two additional hits, Candidate_507 (*LINC00910*) and Candidate_508 (Figure S2g).

**Figure 3.**
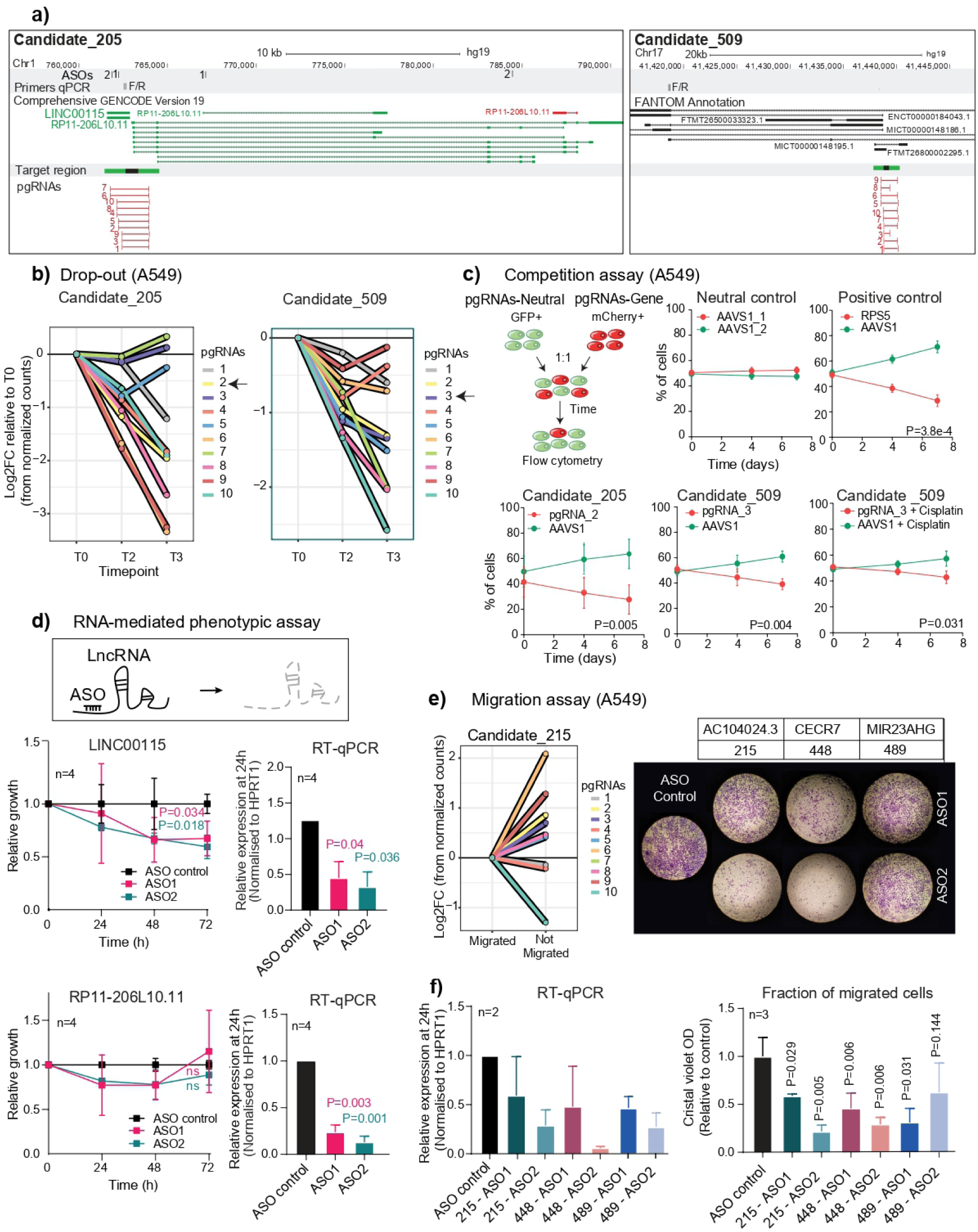
Validation of screen hits shows reproducible phenotypes. a) Candidate_205 (left panel) and Candidate_509 (right panel) loci. Primers and ASO sites are indicated above. The TSS target region and the 10 pgRNAs from the screening library are indicated below. **b)** Normalised pgRNA counts over the course of the drop-out screen in A549. Arrows indicate pgRNAs that were cloned here for validation. **c)** Competition assay. Fluorescently-labelled cells carrying pgRNAs for control *AAVS1* locus (green, GFP) or indicated targets (red, mCherry) were measured by flow cytometry. pgRNAs targeting the ORF of essential ribosomal protein *RPS5* were used as positive control. N=3, error bars indicate standard deviation; statistical significance was estimated by Student’s *t* test at the last timepoint. **d)** ASOs were used to separately target the two lncRNAs sharing the TSS at Candidate_205. For each, two different ASOs were employed (1 and 2) in A549 cells. Upper panels: *LINC00115*; Lower panels: *RP11-206L10.11*. Left: cell population; right: RNA expression measured by RT-qPCR. N=4; error bars indicate standard deviation; significance was estimated by one-tailed Student’s *t* test. **e)** Left: Normalised counts of pgRNAs in migrated and non-migrated cell populations from A549 migration screen. Right: Validation experiments with A549 cell migration across transwell supports over 24 h. Cells were treated with ASOs targeting indicated lncRNAs or a non-targeting control. **f)** Left: RT-qPCR in A549 cells treated with two distinct ASOs each for Candidate_215, 448 and 489. Expression was normalised to a non-targeting ASO. Right: Crystal violet quantification normalised to non-targeting ASO. Data are plotted as mean ± SD from three independent biological replicates. Statistical significance was estimated by one-tailed Student’s *t* test.

To validate the phenotypic effect of these deletions, we tested individual high-scoring pgRNAs (Figure 3b, arrows). Candidate_205 pgRNA efficiently deleted the targeted region (Figure S2h), and while difficulty in designing PCR primers prevented direct testing of deletion by Candidate_509, it effectively decreased RNA levels (Figure S2i). Both pgRNAs yielded potent effects on cell fitness: mCherry+ cells expressing pgRNAs were out-competed by control cells (GFP+, expressing pgRNA for *AAVS1*), with an effect comparable to inactivation of essential ribosomal gene *RPS5* (Figures 3c and S2j). Similar results were observed in a conventional assay (Figure S2k). Furthermore, the pgRNA for Candidate_509 also sensitised cells to cisplatin, consistent with screen results (Figure 3c).

It remained ambiguous which of the two genes overlapping the Candidate_205 region drives these effects. Furthermore, genomic deletion cannot distinguish between a DNA-dependent (for example, enhancer) or RNA-dependent mechanism (mature lncRNA, or its transcription). To address both questions, we used two gene-specific ASOs to target each gene. This clearly implicated *LINC00115*, but not *RP11-206L10*, in driving cell proliferation via an RNA-dependent mechanism (56) (Figure 3d). These results were corroborated in a three-dimensional (3D) spheroid model (Figure S2i).

Similar high validation rates were observed for the migration screens. ASO-knockdown of three hits, Candidate_215 (*AC104024.3*), Candidate_448 (*CECR7*) and Candidate_489 (*MIR23AHG*) resulted in dramatic impairment of A549 migration (Figures 3e and 3f).

In summary, these findings support the ability of CRISPR-deletion screens to identify lncRNA genes that promote cancer hallmarks via RNA-dependent mechanisms.

### Multi-hallmark screen integration for target discovery

We next integrated these data into quantitative and comprehensive map of lncRNAs driving NSCLC hallmarks. To combine diverse screen results while balancing effect size and significance, we created an integrative target prioritisation pipeline (TPP) (Figure 4a). TPP can either be run on all screens for a “pan-hallmark” target ranking, or for individual hallmarks (“hallmark-specific”; see Methods). The pan-hallmark ranking outperforms common integration methods and individual screens in correctly classifying positive and neutral controls (Figure 4b).

**Figure 4.**
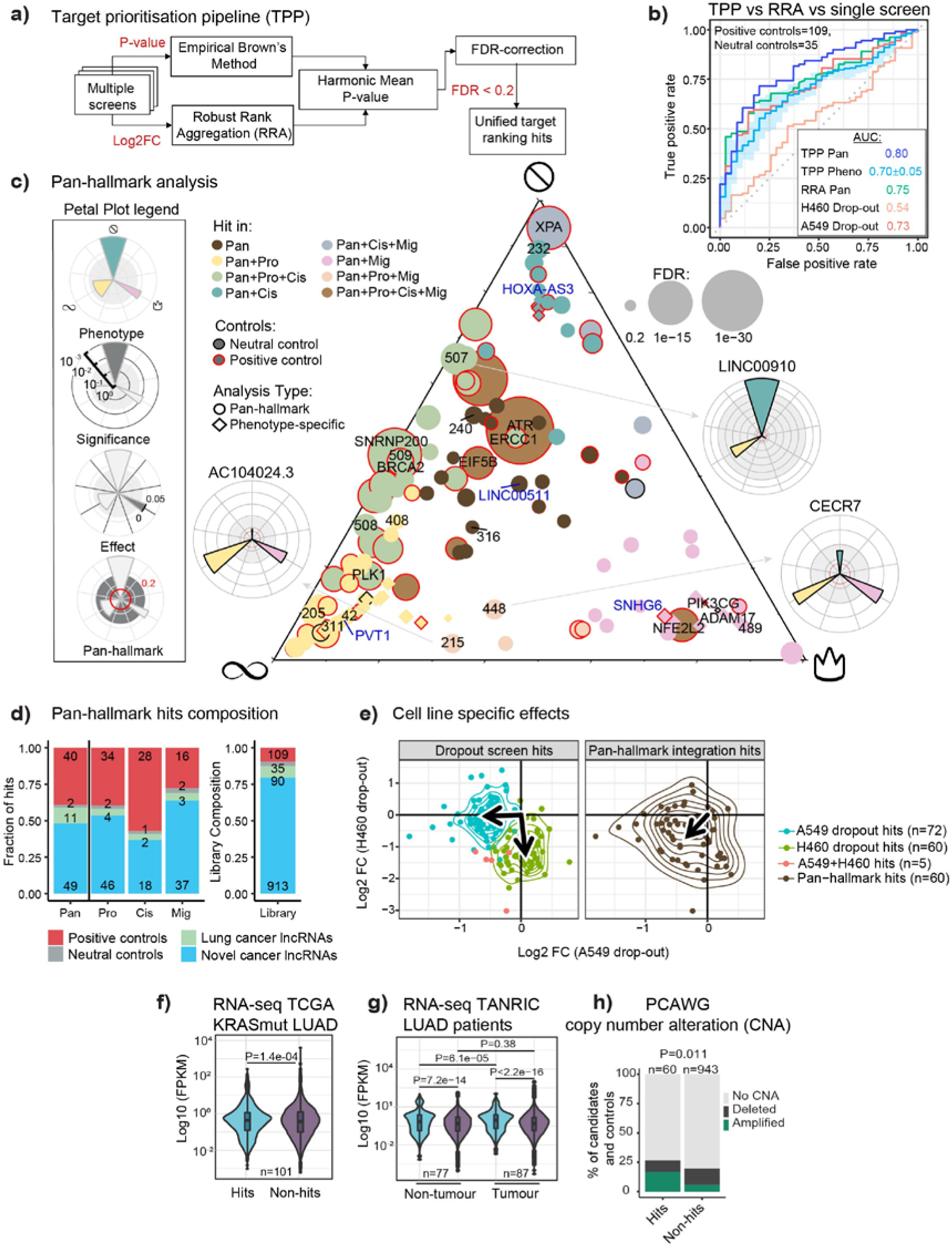
Functional landscape of lncRNAs in NSCLC. a) Target prioritisation pipeline (TPP) integrates multiple screens to generate a unified hit ranking. TPP employs both effect size (Robust Rank Aggregation; RRA) and statistical significance (empirical Brown’s method). **b)** Comparing performance of screens and integration methods. Performance is measured as the area under the ROC curve (AUC) based on correctly recalling/rejecting library controls (positive and neutral controls, respectively). **c)** Ternary plot contains all significant hits (FDR<0.2) in pan-hallmark analysis (circles) or individual hallmarks (diamonds). The three corners represent the hallmarks, indicated by symbols. The proximity to each corner is driven by the TPP significance calculated for each individual hallmark. The candidates selected for further validations and protein-coding controls are indicated by candidate number or gene name, respectively. Selected lncRNAs previously associated with LUAD are labelled in blue. Petal plots display the hallmark contributions of selected lncRNAs. **d)** The number of hits discovered in pan-hallmark and individual hallmark screens. Right: The target composition of the screening library for comparison. “Lung cancer lncRNAs’’ indicate previously-published, functionally-validated lncRNAs in lung cancer(11) **e)** Scatter plot showing the correlation of the drop-out assay hits (left panel) and the pan-hallmark hits (right panel) in H460 and A549. The arrows point to the geometric median of the respective group. **f)** Expression of pan-hallmark hits compared to all other screened lncRNAs (non-hits) in the KRAS^+^ samples from the TCGA LUAD cohort (n=101). Statistical significance was estimated by Welch’s *t*-test, one-tailed. **g**) Expression of pan-hallmark hits compared to non-hits in an independent cohort of LUAD samples and healthy tissues (87 tumour; 77 normal)(60). Statistical significance: pairwise two-tailed Student’s *t*-test. **h**) Pan-cancer recurrent amplifications and deletions, estimated in PCAWG cohort. Statistical significance was estimated by Fisher’s exact test.

The hallmark-specific values provide a signature for each lncRNA in three functional dimensions, in the context of overall confidence defined by pan-hallmark ranking (Figure 4c). The union of hits from both approaches (FDR<0.2) yielded 111 lncRNAs (Figure 4c). Pan-hallmark analysis alone identified altogether 60 lncRNA hits (∼6% of those screened; File S3) (FDR<0.2), of which 49 were not previously linked to NSCLC (Figure 4d). As expected, growth-promoting PCG positive controls and known lung cancer lncRNAs, but not neutral controls, are enriched amongst hits. This is supported by independent Enrichment Score Analysis (Figure S3a). Attesting to their value, hits are significantly enriched for disease-associated lncRNAs (Figure S3b), and consistent with previous reports (57), they are more expressed in healthy tissues and marginally more evolutionarily conserved (Figure S3c). Hallmark-specific integration yielded 96 hits (∼10% of those screened) in at least one hallmark, of which 14 are found in two hallmarks, and none in three (Figure S3d).

Previous CRISPR screens have demonstrated highly background-specific roles in proliferation for cell lines from distinct cancer types(18). However, the degree of similarity between such roles between cell models from a given cancer type is not known. Indeed, comparing lncRNA hits from conventional drop-out analysis in A549 and H460 backgrounds revealed a low degree of correlation (Figure 4e left panel). In contrast, pan-hallmark hits display relatively background-independent activity (Figure 4e right panel), supporting the usefulness of an integrative screening strategy for discovering therapeutic targets.

Cancer-promoting lncRNAs are expected to be upregulated in tumours (58,59). Consistent with this, pan-hallmark hits are significantly higher expressed than non-hits in KRAS^+^ lung tumours (Figure 4f), and are upregulated in tumours compared to adjacent tissue (Figure 4g). Screen hits tend to be amplified, but not depleted, in DNA from tumours (Figures 4h and S3e) and cell lines (Figure S3f).

In summary, integration of diverse screens yields accurate maps of functional lncRNAs that are enriched for meaningful clinical features and display cell background-independent activity.

### RNA therapeutics targeting NSCLC lncRNAs

Multi-hallmark lncRNA maps are a resource of targets for therapeutic ASOs(61–63). We manually selected ten lncRNAs from top-ranked hits, based on criteria of novelty and lack of protein-coding evidence, and henceforth referred to as “Tier 1”. Eight are annotated by GENCODE, and two by either FANTOM CAT or BIGTranscriptome (File S5, Table 1). For each candidate, we designed a series of ASOs and managed to identify at least two independent ASOs with ≥40% knockdown potency (Figure S4a).

Next, we tested ASOs’ phenotypic effects, in terms of proliferation and cisplatin sensitivity (Figure 5a). For five lncRNAs (Tier 2), we observed reproducible loss of cell proliferation with two distinct ASO sequences, indicating on-target activity(64). To check how broadly applicable these effects are, we re-tested the ASOs in two other KRAS^+^ NSCLC cell lines, H460 and H441 (derived from a pericardial effusion metastasis), and observed similar results (Figure 5a). Consistent with their effects on cisplatin sensitivity, Tier 2 genes’ expression is upregulated in response to cisplatin treatment (Figure S4b). Using the expression data from TCGA dataset we noted that Tier 2 lncRNAs are over-expressed in NSCLC tumours, despite this not being a selection criterion (Figure 5b).

**Figure 5.**
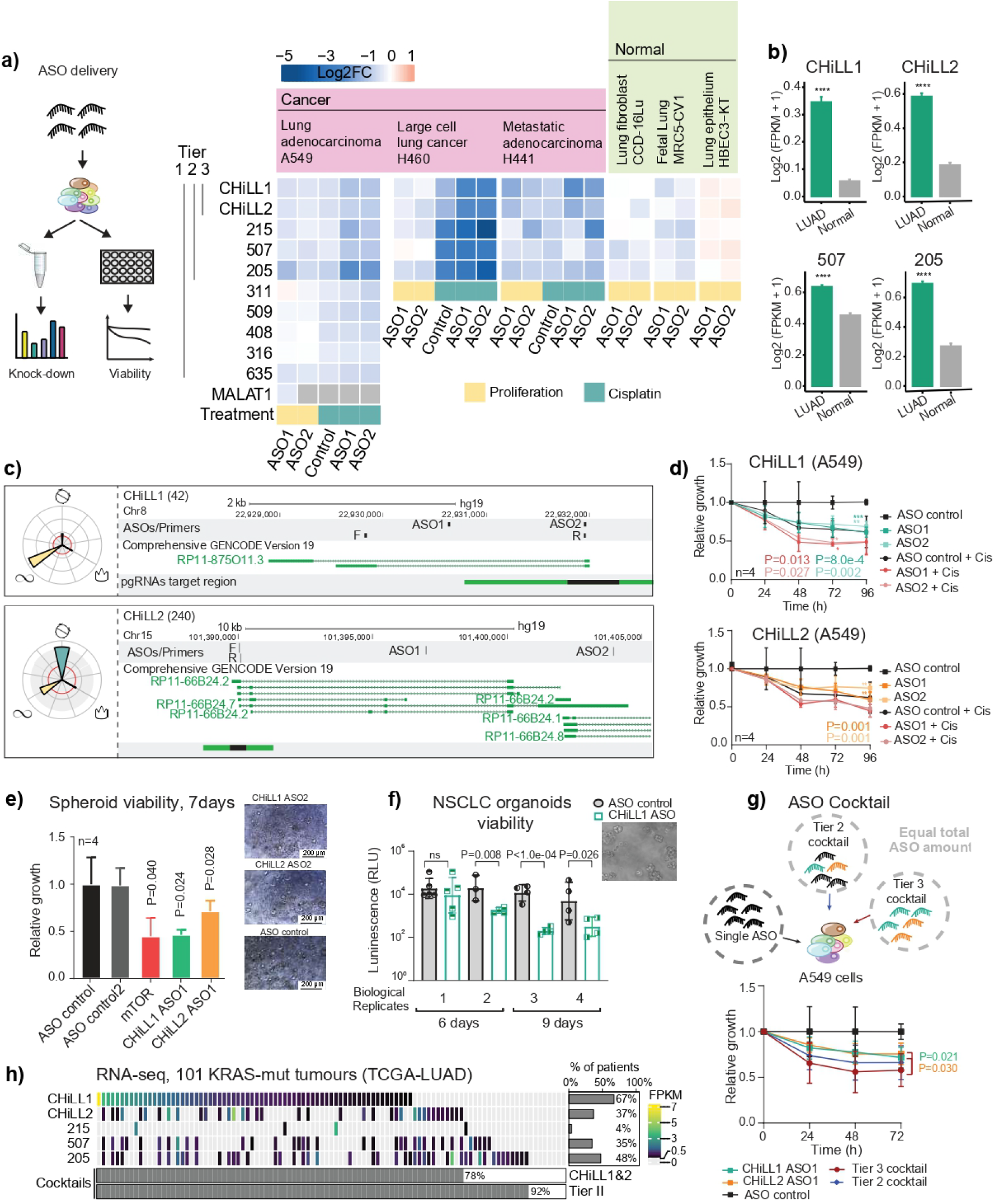
Therapeutic targeting of oncogenic lncRNAs CHiLL1 and CHiLL2. **a)** Left: Experimental workflow to test knockdown efficiency and phenotypic effect of ASO transfection. Right: Summary of all ASO/cell line results. Rows: Targeted lncRNAs; Columns: ASOs and cell lines. Values reflect the mean log2 fold change in viability following ASO transfection, with respect to a control non-targeting ASO, n>2. Numbers to the left indicate library candidate identifiers, and are grouped into Tiers. Each lncRNA is targeted by two independent ASO sequences (1&2, below). MALAT1 lncRNA is used as positive control. “Normal” cells are non-transformed cells of lung origin. **b)** Expression of Tier 2 lncRNAs - CHiLL1, CHiLL2, Candidate_205 (*LINC001150*), Candidate_507 (*LINC00910*) – in TCGA RNA-seq. LUAD: 513 samples, Normal: 59 samples. Statistical significance estimated by *t*-test; **** P<0.0001. Data was not available for Candidate_215 in the TCGA dataset. **c)** Genomic loci encoding CHiLL1 (upper panel) and CHiLL2 (lower panel). **d)** A549 cell growth upon transfection with two independent ASOs. Results are normalised to non-targeting control ASO (n=4 biological replicates; error bars: standard deviation; statistical significance: two-tailed Student’s *t* test). **e)** Left: Viability of H441 spheroid cultures seven days after ASOs transfection (25nM). mTOR ASO was used as positive control (n=4 biological replicates; error bars: standard deviation; one-tailed Student’s *t* test). Right: Representative images (Leica DM IL LED Tissue Culture Microscope). **f**) Viability of BE874 organoids grown from a KRAS^+^ patient-derived xenograft after CHiLL1 ASO transfection (n=4 biological replicates; n>3 technical replicates; error bars: standard deviation; one-tailed Student’s *t* test). g) ASO cocktails. Above: For all experiments, the total amount of ASO did not vary (25 nM). Cocktails were composed of equal proportions of indicated ASOs. Below: A549 cell populations, normalised to non-targeting ASO (n=4 biological replicates; error bars: standard deviation; statistical significance: one-tailed Mann Whitney test). **h**) Expression of Tier 2 lncRNAs in KRAS^+^ LUAD tumours (TCGA). Each cell represents a patient, and is coloured to reflect expression as estimated by RNA-seq (expression defined as >0.5 FPKM). Below, the percentage of patients with at least one lncRNA from the indicated cocktails.

It has been proposed that targeting lncRNAs could cause lower side-effects in healthy tissue, although few studies have tested this(15). We evaluated Tier 2 ASOs’ effects on a panel of non-transformed lung-derived cells: HBEC3-KT, MRC5-CV1 (both immortalised) and CCD-12Lu (primary). These cells displayed diminished or absent response, particularly for the first two candidate lncRNAs (Figure 5a). Consequently, we narrowed our focus to these “Tier 3” lncRNAs: Candidate_42 (*ENSG00000253616*) and Candidate_240 (*ENSG00000272808*), henceforth renamed *Cancer Hallmark in Lung LncRNA* (CHiLL) 1 and 2, respectively. Both have low protein-coding potential (Figures 5c and S4d). Replication experiments confirmed the knockdown potency and phenotypic impacts of both ASOs for each gene (Figures 5d, S4c and S4e). ASOs displayed activity in additional KRAS^+^ and *EGFR*-mutant cell lines, suggesting subtype-independent activity (Figure S4f).

CHiLL1 has, to our knowledge, never previously been implicated in cancer. It is located on Chr8 and consists of two annotated isoforms, sharing the first exon (Figure 5c, upper panel) It is localized upstream and on the same strand of the protein-coding gene *TNFRSF10B*, previously associated with NSCLC(65) although we find no evidence for read through transcription between the two loci (Figures S5a and S5b). Its sequence lacks obvious functional elements (Figure S5c). Supporting its relevance, high expression of CHiLL1 correlates with poor overall survival (Figure S4g).

CHiLL2 (Chr15) comprises four isoforms sharing a common TSS (Figure 5c, lower panel). It is associated with poor prognosis in colon cancer(66), and during preparation of this manuscript, was reported to be an oncogene in gastric cancer(67). It has a likely orthologue in mouse, the uncharacterised Gm44753 (Figure S5f). In contrast to CHiLL1, CHiLL2 exons contain numerous conserved sequences and structures (Figure 6a), and its locus is frequently amplified in cancer genomes (Figure S5d). In TCGA samples, CHiLL2 expression is upregulated in the proximal inflammatory (PI) tumour subtype, which is associated with poorer prognosis (Figures S4h and S4i; PI vs. PP - P=4e-05; PI vs. TRU - P = 0.008).

**Figure 6.**
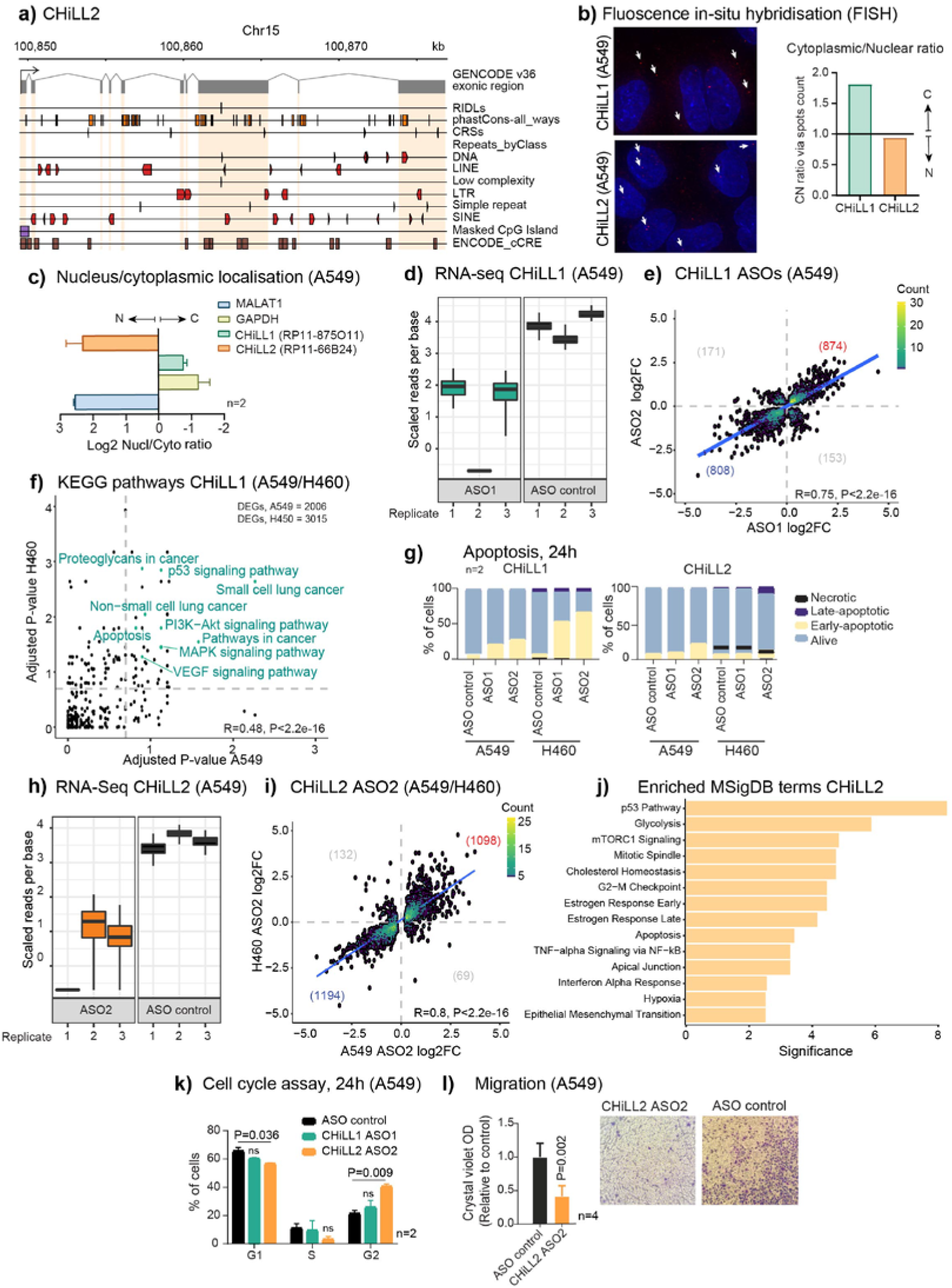
CHiLL1 and CHiLL2 drive distinct but overlapping oncogenic pathways. **a)** Genomic elements in CHiLL2 exons (grey rectangles) and introns. Strand is indicated by element direction, where appropriate. Plot was generated with ezTracks(82). **b)** Confocal microscopy images of RNA-FISH performed with CHiLL1 and CHiLL2 probe sets in A549 cells. Selected lncRNA foci are arrowed. Nuclear/Cytoplasmic (N/C) quantification of CHiLL 1&2. **c)** Ratio of concentrations of indicated RNAs as measured by RT-qPCR in nuclear/cytoplasmic fractions of A549 cells. MALAT1 and GAPDH are used as nuclear and cytosolic controls, respectively. **d)** Expression of CHiLL1 as quantified in the three RNA-seq replicates from A549 RNA. The y-axis represents the normalized expression (counts) per nucleotide and the boxplots show the variance of inference using bootstraps generated by Kallisto. **e)** Fold change in gene expression in response to CHiLL1 knockdown by ASO1 and ASO2 in A549 cells. Statistical significance: Pearson correlation coefficient. Trendline depicts regression line. Numbers indicate genes in each quadrant. f) Statistical enrichment of KEGG pathways amongst DEGs resulting from CHiLL1 knockdown. Genes significantly affected in common by ASO1 and ASO2 were included, for each cell line. Cancer-relevant pathways that are significant for both cell lines are highlighted in green. Statistical significance: Pearson correlation coefficient. **g)** Annexin-V apoptosis assay 24h after transfection with ASOs targeting CHiLL1 (left) and CHiLL2 (right). **h**) As for C, for CHiLL2. **i**) As for D, but comparing knockdown of CHiLL2 with the same ASO in A549 and H460 cells (P=2.2e-16; R=0.8; statistical significance: Pearson correlation coefficient). **j)** MSigDB term enrichment significance for differentially-expressed genes in common between A549 and H460 cells. The most significant terms are shown. y-axis: Adjusted P-value (q-value). **k**) Cell cycle assay results after 24h upon knock-down of CHiLL1 and CHiLL2 in A549. (n=2 biological replicates, error bars: standard deviation; statistical significance: two-tailed Student’s *t* test) **l)** Cell migration across transwell supports in A549 cells. The crystal violet quantification is normalised to non-targeting ASO (n=4 biological replicates, error bars: standard deviation; statistical significance: one-tailed Student’s *t* test).

Three-dimensional (3D) *in vitro* models represent a more faithful tumour model compared to monolayer cultures(68,69). We delivered CHILL1&2 ASOs to spheroid cultures of H441 cells, and observed a reduction in viability approaching that of the positive control, mTOR (Figures 5e and S4j).

Organoids derived from patient-derived xenografts (PDX) recapitulate the therapy response of individual patients(70). Delivery of CHiLL1 ASOs resulted in significant reduction in cell viability of the KRAS^+^ human NSCLC organoid BE874 (Figure 5f).

We were curious whether simultaneous targeting of two or more distinct vulnerabilities, via “cocktails” of ASOs, might offer synergistic benefits. Indeed, a 50:50 cocktail of CHiLL1 / CHiLL2 ASOs (Tier 3 cocktail) displayed a greater effect on cell viability, compared to an equal dose of either ASO alone (Figures 5g and S4k). A five-ASO cocktail for Tier 2 lncRNAs yielded a similar benefit (Figure 5g). Interestingly, cocktails resulted in no additional toxicity for non-cancerous cells (Figure S4l).

Finally, we asked what fraction of patients might benefit from ASO treatment. RNA-sequencing (RNA-seq) data for TCGA tumours indicates that 78% express at least one of CHiLL1 and CHiLL2 and might be treated by the Tier 3 cocktail, rising to 92% for Tier 2.

Together these results demonstrate that Tier 3 lncRNAs can be targeted by potent and low-toxicity ASOs, which may be beneficial alone or in combination for the majority of NSCLC cases (Figure 5h).

### CHiLL1&2 ASOs regulated cancer hallmarks via distinct modes of action

We next investigated the modes of action linking CHiLL1&2 to NSCLC hallmarks. Subcellular localisation yields important mechanistic clues for lncRNAs (71). Surprisingly, despite their similar oncogenic roles, fluorescence *in situ* hybridisation (FISH) revealed contrasting localisation patterns: CHiLL1 is located principally in the cytoplasm, and CHiLL2 in the nucleus (Figure 6b). Specificity was validated by knockdown (Figure S5e), and results were further corroborated by cell fractionation (Figure 6c).

To gain more detailed mechanistic insights, we used molecular phenotyping by RNA-seq to quantify the transcriptome of A549 cells perturbed by CHiLL1 ASOs(72). CHiLL1 expression in control cells was 8.3 TPM (transcripts per million), equivalent to ∼4 molecules per cell and consistent with FISH(73). RNA-seq confirmed ASO knockdown efficiency (Figures 6d and S6a), and resulting transcriptome changes were highly correlated between ASOs, indicating that the majority of effects arise via on-target perturbation of CHiLL1 (Figure 6e). Similar correlation was observed in H460 cells (Figure S6b). The generality of CHiLL1-dependent expression changes was further confirmed by correlated expression changes between A549 and H460 (Figure S6c).

We explored perturbed genes by enrichment analysis. Defining high-confidence target gene subsets from the intersection of both ASOs, we identified enriched KEGG terms(74) (Figures 6f and S6d). These underscored disease relevance (e.g. “Non-small cell lung cancer”), and also implicated potential mechanistic pathways (MAPK, PI3K-Akt), and the high degree of concordance between the two cell backgrounds again supported the generality of CHiLL1 effects across KRAS^+^ NSCLC cells.

Similar analysis of the Molecular Signatures Database (MSigDB)(75) implicated p53 and mTORC1 signalling (Figure S6e). Numerous transcription factor binding sites (TFBS) are enriched in changing genes, including *ZBTB7A* (in both A549 and H460)(76,77) (Figure S6f). Interestingly, *ZBTB7A* has been reported as both oncogene and tumour suppressor that regulates processes including apoptosis and glycolysis (78–81).

Amongst the cell-type independent enriched KEGG pathways was “Apoptosis” (Figure 6f). Supporting this, knockdown of CHiLL1, but not CHiLL2, resulted in a significant increase of early apoptotic cells (Figure 6g).

Turning to CHiLL2 (mean expression 2.5 TPM, ∼1 copies per cell), we observed effective knockdown and cell-type-independent effects using a single ASO (Figures 6h, 6i, and S6g). Gene enrichment analysis revealed partially overlapping terms with CHiLL1 (including p53 pathway, cholesterol homeostasis and epithelial to mesenchymal transition). Amongst the CHiLL2-specific terms, we noticed several related to cell cycle progression, including “G2−M checkpoint” (Figure 6j). Indeed, knockdown of CHiLL2, but not CHiLL1, led to an increase of cells in G2-phase (Figure 6k). CHiLL2 targets are also enriched for migration genes, and knockdown led to impaired cell migratory capability (Figure 6l).

Interestingly, we notice that CHiLL2 knockdown resulted in a significant upregulation of CHiLL1, but not vice versa, (Figure S6h), explaining the additive effect on cell viability observed with the CHiLL1 / CHiLL2 ASO cocktail (Figure 5g).

Together, these data establish that CHiLL1&2 promote cancer hallmarks via widespread, non-overlapping downstream gene networks, and support the on-target basis for ASO activity.

## Discussion

We have mapped the functional lncRNA landscape across hallmarks and cell backgrounds in the most common KRAS^+^ subtype of NSCLC. This led us to promising therapeutic targets with pleiotropic roles across a range of 2- and 3-dimensional models from primary and metastatic tumours.

Pooled CRISPR screening is emerging as a foundation for therapeutic target identification, thanks to its practicality and versatility (83). It avoids the investment and expertise required for arrayed screening(84), and delivers improved perturbation and on-target rates compared to shRNA(35). By directly identifying lncRNAs via their cell-level function, it represents a welcome addition to widely-used, indirect evidence like survival, mutation, differential expression or evolutionary conservation(85,86). It was encouraging to observe concordance between these signatures and screen hits. Nonetheless, key barriers to entry remain, notably the lack of available screening libraries, making the libDECKO-NSCLC1 library a valuable resource for future discovery.

Through its integrative strategy, this work sets new standards. Previous studies typically screened a single cell line with a single hallmark (often proliferation) and in a single format (often drop-out)(18,20,87). Internal benchmarking here highlighted the risks of this approach, in identifying hits whose activity is specific to that cell background alone. We mitigated this with parallel screens in distinct but matched cell backgrounds, and resulting hits could be validated in several additional models and mutational subtypes. To identify pleiotropic hits, we performed parallel screens in multiple phenotypic dimensions and both positive and negative formats. The resulting challenge of integrating diverse screen data was solved by developing the simple yet robust TPP pipeline that balances effect size and significance. Overall, the combination of multiple screens with TPP yielded improved performance compared to conventional approaches.

The result is a unique functional panorama of lncRNAs in a single cancer type. Given the paucity of global-level functional lncRNA maps(18,88), this dataset represents an invaluable resource for understanding both the basic biology of lncRNAs and their roles in cancer. Overall, our data implicate approximately 6% of lncRNAs with cell-level functions, comparable to Liu’s and Zhu’s estimates(18,40). The conceptual and experimental configuration will have a broad application for other diseases and biological systems.

This resource enabled us to narrow down ten lncRNAs for ASO development, from which we identified a pair of oncogenic lncRNAs, CHiLL 1 &2 with particularly promising characteristics as therapeutic targets. Firstly, replication experiments using distinct ASO sequences and in different cell backgrounds strong suggested that observed phenotypic and molecular effects occur on-target (via the intended lncRNA)(89). Second, ASOs were effective in both monolayer and three-dimensional KRAS^+^ NSCLC backgrounds, in addition to several EGFR-mutant cell lines, raising hope for a more general utility. Future efforts will be required to further refine ASO sequences and chemistry, and to effectively deliver them *in vivo*. Third, those phenotypic effects were diminished in non-transformed cells, pointing to reduced non-specific toxicity *in vivo*. This raises hopes for reduced toxicity in healthy cells, allowing not only higher doses, but also combination therapies to suppress therapy resistance(90). Finally, CHiLL1 or 2 are detected in the majority of KRAS^+^ NSCLC tumours, suggesting the majority of patients might benefit from eventual treatment.

Mechanistically, the concordance of CRISPR and ASO phenotypes indicates that both genes act via an RNA transcript, or at least the production thereof(56,91). Both have profound effects on the cellular transcriptome, affecting hundreds of target genes that converge on many shared, oncogenic pathways, which yielded experimentally verifiable predictions. Interestingly, however, these effects are mediated by very different immediate molecular mechanisms, as evidenced by their distinct subcellular localisation, non-overlapping target genes, and the fact that mixing their ASOs yielded greater than additive effects.

We have shown how ASO cocktails boosted efficacy without increasing toxicity, compared to equal doses of single ASOs. Our findings open the possibility of using either fixed cocktails or cocktails tailored specifically to a patient’s tumour transcriptome, for potent, enduring, low-toxicity and personalised cancer treatment.

Nonetheless several issues remain to be addressed in future. It is likely that we overlooked many valuable lncRNA targets, due to the relative inefficiency of CRISPR-deletion as a perturbation compared to ORF mutation(92), and also due to the ongoing incompleteness of lncRNA annotation(93). We here screened in two monolayer cell backgrounds, however future screens should be performed in parallel across the widest panel of mutationally-matched 2- and 3-dimensional models(94). Finally, our screens discovered scores of lncRNAs that could not be followed up here and hopefully will provide a fertile source of new targets in the future.

## Additional information

## Acknowledgements

We thank members of the GOLD Lab, including Joana Carlevaro-Fita, Núria Bosch-Guiteras, Michela Coan, Antonio Tarruell and Tina Uroda for insightful discussions. We also thank Roderic Guigó (CRG Barcelona) for insightful discussion. Thomas Marti and Renwang Peng (DMBR) generously donated cell lines and advice on functional assays. We thank Basak Ginsbourger (DBMR) for administrative support, and Willy Hofstetter and Patrick Furer (DBMR) for logistical support. All computation was performed on the Bern Interfaculty Bioinformatics Unit computing cluster maintained by Rémy Bruggmann and Pierre Berthier.

## Author contributions

R.E., T.P., and R.J. conceived and designed the experiment procedure and performed data analysis and its interpretation. D.F.M developed the TPP pipeline and performed most of the bioinformatic analysis C.P. Designed the ‘libDECKO-NSCLC1’ with the help of S.H. P.C. Analyzed the RNA-seq data. S.F. and M.R. Established the patients-derived organoids. P.S., AK, G.B. and J.S. generated the stably-expressing Cas9 cell line lines and helped with the validation. E.S.W., I.M., and P.C.M. validated the results in additional cell lines. D.H. performed the analysis on the FISH images. L.Z. and X.W. performed the survival analysis from TCGA data. A.V. provided the set of known cancer genes in NSCLC. A.A., H.A.G., P.P.M. and A.L. contributed to the bioinformatic analysis. I.C. prepared the libraries for the NGS sequencing. S. H., C.R., and A.O. provided key inputs and tools. T.P., R.E. and R. J. wrote the manuscript with input from all the authors.

## Availability of data and material

The data generated during this study will be available within few days in the Gene Expression Omnibus repository (GEO).

## Abbreviations

ASO: Antisense oligonucleotide
CRISPR: Clustered regularly interspaced short palindromic repeats
CHiLL: Cancer Hallmark in Lung LncRNA
DECKO: Dual excision CRISPR knock-out
FDR: False Discovery Rate
FISH: Fluorescence in situ hybridisation
FPKM: Fragments per kilobase of exon per million mapped fragments
KEGG: Kyoto Encyclopedia of Genes and Genomes
KO: Knock-Out
KRAS: Kirsten rat sarcoma virus
LNA: Locked Nucleic Acid
LncRNA: Long non-coding RNA
MOI: Multiplicity of infection
NGS: Next-Generation Sequencing
NSCLC: Non small cell lung cancer
ORF: Open Reading Frame
PCR: Polymerase Chain Reaction
pgRNA: paired guide RNAs
qPCR: quantitative Polymerase Chain Reaction
RNA-seq: RNA-sequencing
RRA: Robust Rank Aggregation
TCGA: The Cancer Genome Atlas
TFBS: Transcription factor binding sites
TPM: Transcripts per million
TPP: Target prioritisation pipeline
TSS: Transcriptional Start Site

## Methods

### Cell lines and culture

HEK293T, A549, H460, H441, CCD-16Lu cell lines were a kind gift by the groups of Adrian Ochsenbein and Renwang Peng (University Hospital of Bern). MRC-5 cells were provided by the group of Ronald Dijkmanthe (Institute of Virology and Immunology, University of Bern). HBEC3-KT bronchial epithelial human cells were purchased from the American Type Culture Collection (ATCC; http://www.atcc.org). All the cell lines were authenticated using Short Tandem Repeat (STR) profiling (Microsynth Cell Line Typing) and tested negative for mycoplasma contamination.

A549 and HEK293T cells were maintained DMEM, MRC-5 in EMEM, NCI-H460, H441, and CCD-16Lu in RPMI-1640 medium, all supplemented with 10% Fetal Bovine Serum, 1% L-Glutamine, 1% Penicillin-Streptomycin. HBEC3-KT were maintained in Airway Epithelial Cell Basal Medium (ATCC®, cat. no. PCS-300-030) supplemented with Bronchial Epithelial Cell Growth Kit (ATCC®, cat. no. PCS-300-040).

All cells were passaged every 2−3 days and maintained at 37°C in a humid atmosphere with 5% CO2.

### Lentiviral infection and stable cell line production

The plasmids used in this paper are listed in File S4. Lentivirus production was carried out by co-transfecting HEK293T cells with 12.5 μg of Cas9 plasmid with blasticidin resistance (Addgene, cat. no. 52962), 7.5 μg psPAX2 plasmid and 4 μg the packaging pVsVg plasmids, using Lipofectamine2000. 24h before the transfection, 2.5e6 HEK293T cells were seeded in a 10 cm dish coated with Poly-L-Lysine (Sigma, cat. no. P4832) (diluted 1:5 in 1X PBS). The supernatant containing viral particles was harvested 24h, 48h and 72h after transfection. Viral particles were then concentrated 100-fold by adding 1 volume of cold PEG-it Virus Precipitation Solution (BioCat, cat. no. LV810A-1-SBI) to every four volumes of supernatant. After 12h at 4°C, the supernatant/PEG-it mixture was centrifuged at 1,500 × g for 30min at 4°C, resuspended in 1X PBS, and stored at −80°C till use.

For the generation of stable Cas9-expressing cell lines, A549 and H460 were incubated for 24h with culture medium containing concentrated viral preparation carrying pLentiCas9-T2A-BFP and 8 μg/ml Polybrene. Infected cells were selected for at least five days with blasticidin (8 μg/mL) and then were FACS-sorted two times, so as to have at least 60% BFP-positive cells.

### Design and cloning of DECKO plasmids and lentiviral production

For the design and cloning of DECKO plasmids, we used our previously-described protocol(23,95) (http://crispeta.crg.eu/).

To produce lentivirus carrying the pDECKO plasmid, we followed the same protocol. After infection with pDECKO plasmid-carrying viruses, cells were selected with puromycin (μg/mL) for at least three days.

### Library design

We downloaded GTF-format annotations from the following sources: i) GENCODE annotation release 19 (GRCh37) from gencodegenes.org; ii) BIGTranscriptome annotation(28) from http://big.hanyang.ac.kr/CASOL/ l; iii) FANTOM CAT(10). We also generated a novel transcriptome assembly of A549 RNA-seq(30,31) using StringTie(96), version 1.3.

All lncRNAs were filtered thus: First, those with transcription start sites (TSS) <2kb from any protein-coding gene exon were removed. Second, expression was calculated with RSEM v1.3 (97), and transcripts with FPKM <0.1 were removed. Remaining TSS within 300 bp were clustered into a single TSS. TSS were intersected with ENCODE evidence source specific to A549 cells: CAGE, DNAse I hypersensitivity sites and ChromHMM marks: Active TSS, Flanking TSS, Promoter Downstream TSS, Flanking TSS Downstream, Genic enhancer1, Genic enhancer2, Active Enhancer 1, Active Enhancer 2, Weak Enhancer and Bivalent-Poised TSS (30,31,98). Candidates were prioritized by the number of evidence sources.

We designed neutral control pgRNAs in genomic regions not expected to affect cell phenotype. We retrieved 10 regions in the AAVS1 gene loci from the publication of Zhu and colleagues (99). To this set we added a set of 65 randomly selected intergenic regions (>10 kb distant from nearest gene annotation) and 25 intronic regions (for introns >5 kb in length). Moreover, 53 positive (promoting cell growth) and 50 negative (opposing cell growth) protein-coding gene (PCG) controls, with known roles promoting/opposing cancer cell growth and cisplatin resistance were added. These were manually selected from literature and retrieved from the paper of Zhu and colleagues(99). The complete list of genes contained in the library is available in File S1.

10 unique pgRNAs were generated for each candidate region with CRISPETa(95) using the following parameters: -eu 0 -ed 0 -du 1000 -dd 1000 -si 0.2 -t 0,0,0,x,x -v 0.4 -c DECKO. For the candidates where <10 pgRNAs could be identified, the parameters were subsequently loosened until 10 were reached: in the second round one off-target with 3 mismatches was allowed; in the third round the designs region was repeatedly increased in size (summary in File S5, Table 2).

The final library design comprised 12,000 unique sequences of length 165/166 bp, with overhangs compatible with cloning into the pDECKO plasmid(23,95). The median distance between the pgRNAs is shown in Figure S1d.

### Library cloning

Library was synthesized as single stranded oligonucleotides by Twist Bioscience (USA), and upon arrival resuspended in nuclease-free low Tris-EDTA (TE) buffer (10 mM Tris-HCl, pH 8.0 and 0.1 mM EDTA) to a final concentration of 10 ng/μl. This was PCR amplified using the Oligo-Fw: 5’-ATCTTGTGGAAAGGACGAAA-3’ and Oligo-Rev: GCCTTATTTTAACTTGCTATTTC (PCR mix listed in File S5, Table 3) with the following conditions: 95°C x 1min; 10 cycles of (95°C x 1 min, 53°C x 20 sec, 72°C x 1 min); 72°C x 10 min. The amplification product was purified using the QIAquick PCR Purification Kit (Qiagen, cat. no. 28104) according to the manufacturer’s instructions. The correct amplicon size was checked on a 2% agarose gel at 100V for 40 min.

The steps of cloning follow the low-throughput protocol described in (100). In the first step, the pDECKO_mCherry plasmid was digested following the conditions listed in File S5, Table 4. The amplified library was inserted into the digested plasmid, using Gibson Assembly mix (obtained from ‘Biomolecular Screening & Protein Technologies’ Unit at CRG, Barcelona) at 50 °C for 1 h (200ng of pDECKO_mCherry plasmid, 20ng amplified library, H2O up to 10 μl, 10 μl of Gibson mix 10 μl). 1 μl of the Gibson reaction was delivered to 25 μl of electrocompetent EnduraTM cells (Lucigen, cat. no. 60242-2) using Gene Pulser®/MicroPulser™ Electroporation Cuvettes, 0.1 cm gap (Biorad, cat. no. 16520891). The library coverage of 66.7X was estimated by counting the number of obtained bacterial colonies divided by the total number of different sequences in the designed library (12,000). The intermediate plasmid obtained in this step contains the pgRNA variable sequences, but still lacks the constant part of the first sgRNA and the H1 promoter (Figure S1b)(23).

In the second step of cloning, the intermediate plasmid was digested by BmsbI enzyme (ThermoFisher, cat. no ER0451). After purification, the constant insert was assembled by ligation, by using PAGE purified and 5’ phosphorylated long oligos (File S5, Table 5), as explained in(100). Afterwards, 5 μl of the ligation product was transformed and used for the electroporation of electrocompetent Endura™ cells as described above. Clones were tested by colony PCR and by Sanger sequencing using primer sequences found in File S5, Table 6. The PCR conditions are listed in File S5, Table 7. The overall library quality was evaluated by NGS sequencing. Briefly, the plasmid containing the pgRNAs was amplified by PCR (primers listed in File S5, Table 8), purified using Agencourt AMPure XP beads (Beckman Coulter, cat. no. A63880), according to the manufacturer’s protocol. The purified product was sequenced by Illumina at a depth of 20M PE125 reads. The reads were aligned to the pgRNAs library and the read distribution of each pgRNA was determined using the Ineq package in R (version 3.5.3) to calculate both the Lorenz-curve and Gini-coefficient (Figure S1c).

### Lentiviral titer calculation and lentiviral infection

To achieve the desired multiplicity of infection (MOI) of 0.3–0.4, a titration experiment in A549 and H460 cells was performed. 2e6 cells were plated in each well of a 12-well plate and supplemented with 8 μg/ml polybrene. Each well was treated with virus ranging from 2.5 and 50 μl and transduced via spin-infection as previously described (101). After centrifugation, the media was replaced with complete fresh media without polybrene and incubated overnight. The following day, cells were counted and each well was split in two equal aliquots, of which one was treated with 2 μg/ml puromycin. After 72 h, the MOI was calculated by dividing the number of surviving cells in the puromycin well, by the number in the puromycin-free well. The MOI of 0.3 was used for all screening experiments. For large-scale screens, 120M cells were seeded in 12-well plates with a density of 2M per well for spin-infection. The following day, cells were pooled together and fresh puromycin-containing (2 μg/ml) medium was added. Puromycin selection was maintained for six days until phenotypic screens began.

### CRISPR screens

One week after infection (Timepoint 0 or T0), cells were counted and the reference sample was collected (T0, 16M cells corresponding to a library coverage >1,000x). For all screens, cells were cultured in 150 mm culture-treated dishes and passaged every 2-3 days.

#### Proliferation

Drop-out screens: at T0 16M of cells were plated and passaged so as to maintain a coverage >1,000X (defined as the number of cells divided by the number of unique library sequences). Cells were harvested at 14 and 21 days for gDNA extraction. CFSE screens: At T7, 16M cells were seeded and starved for 24 h with media lacking FBS. Then cells were stained using CellTrace™ CFSE Cell Proliferation Kit (ThermoFisher, cat. no. C34570) following the manufacturer’s instructions. One aliquot of stained cells was immediately analyzed by flow cytometry, while the rest were plated with normal media. Five days later (T5), cells were sorted into two populations: 20% brightest (slow-growing) and 20% least bright (fast-growing). The two populations were plated separately and, five days later (T10) subjected to another round of staining and sorting.

#### Cisplatin screen

Optimal cisplatin working concentrations were established via dose response (Figure S2d) and cell doubling time (Figure S2e). In the dose response, 3,000 A549 and H460 cells were plated in 96-well plates and treated with a range of cisplatin concentrations. After 72 h, CellTiter-Glo 2.0 (Promega, cat. no. G9242) was added to the media (1:1), and luminescence was recorded. For the cell doubling time, 1M cells were plated in 10 cm plates. Different cisplatin concentrations were added at indicated concentrations, and living cells counted every 2-3 days up to 14 days. Cisplatin survival screen: 48M and 96M cells were plated at T0 and treated with 6.5 μM and 25 μM of Cisplatin for A549 cells and 2 μM and 10 μM for H460 cells, corresponding to IC30 and IC80, respectively. Cell pellets were collected after 14 and 21 days. The death screen was carried out as follows: 144M cells were seeded and treated with cisplatin at 2 μM and 1 μM (IC20) for A549 and H460, respectively (Figure S2d). Every 24 h, for five days, floating (dead) cells were collected and pooled together for gDNA extraction.

#### Migration screen

To test the optimal conditions, the following set-up experiment was performed. 0.5 M A549 cells/well were seeded in 5 Boyden chambers (Corning, PC Membrane, 8.0µm, 6.5mm, cat. no. 3422-COR). Each migration assay was stopped at a different timepoint (ranging from 5 h up to 48 h; Figure S2g). 48 h was selected as timepoint for the following experiment. At T0 infected cells (∼16M) were divided and seeded in the upper part of 32 transwell inserts (0.5 M cells/transwell). The upper part of transwell inserts was filled with media lacking FBS, the lower part with media containing 10% FBS. After 48 h cells in the upper part of the chamber (impaired migration) and lower part (accelerated migration) (Figure 2c) were trypsinized and plated separately for 48 h, after this time, cells were counted and collected for gDNA extraction. Control cells that did not undergo the migration assay were harvested at the same time as a reference population.

### Genomic DNA preparation and sequencing

Genomic DNA (gDNA) was isolated using the Blood & Cell Culture DNA Midi (5e6–3e07 cells) (Qiagen, cat. no. 13343), or Mini (<5e6 cells) Kits (Qiagen, cat. no. 13323) as per the manufacturer’s instructions. The gDNA concentrations were quantified by Nanodrop.

For PCR amplification, gDNA was divided into 100 μl reactions such that each well had at most 4 μg of gDNA. Each well consisted of 66.5 μl gDNA plus water, 23.5 μl PCR master mix (20 μl Buffer 5X, 2 μl dNTPs 10 μM, 1.5 μl GoTaq; Promega, cat. no. M3001), and 5 μl of Forward universal primer, and 5 μl of a uniquely barcoded P7 primer (both stock at 10 μM concentration). PCR cycling conditions: an initial 2 min at 95 °C; followed by 30 s at 95 °C, 40 s at 60 °C, 1 min at 72 °C, for 22 cycles; and a final 5 min extension at 72 °C. NGS primers are listed in File S5, Table 9 and Table 10. PCR products were purified with Agencourt AMPure XP SPRI beads according to manufacturer’s instructions (Beckman Coulter, cat. no. A63880). Purified PCR products were quantified using the Qubit™ dsDNA HS Assay Kit (ThermoFisher, cat. no. Q32854). Samples were sequenced on a HiSeq2000 (Illumina) with paired-end 150 bp reads at coverage of 40M reads/sample.

### Screen hit identification and prioritisation

The raw sequencing reads from individual screens were analyzed by using CASPR(102). After the mapping step, the obtained counts per million (cpm) for each pgRNA were filtered to remove sequences with 3>cpm>666. Low scoring guides were removed by GuideScan(103), and a batch effect correction was applied using MageckFlute(104). After all the corrections, the table count was provided to CASPR to calculate log2-Fold Change and FDR corrected P-values at a target level.

To integrate multiple screens an integrative target prioritisation pipeline (TPP) was designed, applying two different approaches in parallel: the Robust Rank Aggregation (RRA)(105) to compute a ranking based on the effect size (CASPR log2FC) across screens; and an empirical adaptation of Brown’s method (EBM)(106) to combine the significance values (CASPR P-value) of each candidate across screens. The RRA-scores were converted to exact P-values using the rho-score correction from the same R package. Subsequently, the harmonic mean P-value (HMP)(107) was calculated using the two significance scores from RRA and EBM. These P-values were corrected for multiple hypothesis testing using the Benjamini & Hochberg method, and a cutoff of FDR<0.2 was used to define hits. The code is available at https://github.com/RescueGum/TargetPP.

Enrichment scores and nominal P-values (GSEA simulation, n=10,000) of positive and neutral control genes were used as indication for the quality of the ranking, as well as fraction of detected genes previously linked to lung cancer(11). Positive and neutral control genes were also used as “true positives/false negatives” and “false positives/true negatives” respectively to calculate ROC curves and associated statistical metrics.

### Public RNA-sequencing data

101 KRAS^+^ LUAD RNA-seq samples were downloaded from TCGA (gdc.cancer.gov), applying the following filters: Adenocarcinoma - not treated - KRAS mutated, and the expression of target genes was estimated using HTSeq(108). Another independent cohort of LUAD RNA-seq ex-vivo data, containing 87 tumour and 77 adjacent normal tissue samples, was obtained from the TANRIC(60,109).

### PCR amplification from genomic DNA

gDNA was extracted with GeneJET Genomic DNA Purification Kit (ThermoFisher, cat. no. K0702) from pDECKO-transduced A549-Cas9-expressing cells. The PCR was done with primers flanking the deleted region (File S5, Table 11) as shown in Figure 1F, using the Phusion™ High-Fidelity DNA Polymerase (2 U/µl) (ThermoFisher, cat. no. F-530S). The product was run on a 1% agarose gel.

### Competition assay

A549 cells were infected with DECKO lentiviruses expressing fluorescent proteins. Viruses expressing control pgRNAs targeting *AAVS1* also expressed GFP protein (pgRNAs-AASV1-GFP+), while the pgRNAs targeting candidate lncRNAs expressed mCherry. After infection, and seven days of puromycin (2 μg/ml) selection, GFP and mCherry cells were mixed 1:1 in a six-well plate (150,000 cells). Cell counts were analyzed by LSR II SORP instrument (BD Biosciences) and analyzed by FlowJo software (Treestar).

### Patient-derived xenograft organoids

The KRAS^+^ patient-derived organoid BE874 was derived in the following way. Small pieces (∼1 – 2 cm^3^) of lung cancer tissue (provided by the Institute of Pathology, University of Bern) were taken from the surgically resected lung cancer specimen with patients’ informed consent. Parts of the sample (pieces of around 5 mm) were separated and implanted subcutaneously into the flanks of 6 weeks old NOD.Cg-PrkdcscidIl2rgtm1Wjl/SzJ (NSG) mice (purchased from Charles River Laboratories) for cancer engraftment(110). After successful engraftment, tumour bearing mice were euthanized and tumours were resected. Single cells were isolated through mechanical and enzymatic tissue disruption for generation of BE874 organoids. Genotyping of BE874 organoids was performed at the Institute of Pathology, University of Bern, using *KRAS* targeted sanger sequencing. KRAS c.34G>T (p.Gly12Cys) mutation was detected in both BE874 organoids and the corresponding primary cancer.

NSG mice were housed under specific pathogen-free conditions in isolated ventilated cages on a regular 12-hour/12-hour cycle of light and dark. Mice were fed ad libitum, and were regularly monitored for pathogens. Mouse experiments were licensed by the Canton of Bern and were performed in compliance with Swiss Federal legislation.

### Antisense oligonucleotides

Locked nucleic acid ASOs were designed using the Qiagen custom LNA oligonucleotides designer (www.qiagen.com). Per each target, we designed from 3 to 5 different ASOs. The day of the transfection, 300,000 cells were counted and plated on a 6-well plate. ASOs were transfected into the cells still in suspension, using Lipofectamine3000 (ThermoFisher, cat. no. L3000015) with final 25 nM in 2 ml media for A549, H460, NCI-H441, and MRC5-CV1 and 10 nM in 2 ml for HBEC3-KT and CCD-16LU, following the manufacturer’s instructions.

For cocktail experiments the final concentration of the ASOs mix was kept at 25 nM. The media was refreshed 24h post transfection and cells were harvested to check the efficiency of gene knockdown or sub-cultured for cell viability experiments. The ASOs target sequences are listed in File S5, Table 1. We checked ASOs penetration in cells by means of the 5′-FAM-labeled control ASO A provided by Qiagen (Figure S5f).

### 2D cell viability assay

Cell viability assay was carried out in 2D cell lines by using CellTiter-Glo 2.0 (Promega, cat. no. G9242). The assays were performed according to the corresponding manufacturer’s protocol. 24h after the transfection, A549, H460, NCI-H441, H1975, H157, WVU-Ma-0005A, H820 and H1650 cells were harvested, counted and 3,500 cells/well were seeded in triplicate in 96 well plates. For Mrc5-SV1, HBEC3-KT and CCD-16LU 3,000, 3,500, and 1,000 cells/well were seeded, respectively. The number of viable cells was estimated after 24, 48, 72, 96 and/or 144h. The day of the measurement, a mix of 1:1 media and CellTiter-Glo was added to the plates and the luminescence was recorded with Tecan Infinite® 200 Pro. Student’s *t* test was used to evaluate significance (P<0.05).

### 3D cell viability assay

NCI-H441 cells were detached, counted, and 200,000 cells were plated in 24 well plates. The ASO-Lipofectamine3000 mix was delivered to the cells in suspension as described above. After 24 hours, the cells are detached, counted and seeded onto 96-well Black/Clear Round Bottom Ultra-Low Attachment Surface Spheroid Microplate (Corning, cat. no. 4520) in 20 μl domes of Matrigel® Matrix GFR, LDEV-free (Corning, cat. no. 356231) and RPMI-1640 growth medium (1:1) with a density of 20,000 cells per dome. Matrigel containing the cells was allowed to solidify for an hour in the incubator at 37 °C before adding DMEM-F12 (Sigma, cat. no. D6421) media on top of the wells (40 μl and 80 μl for the wells intended to the first and second timepoint, respectively. The spheroids were allowed to grow in the incubator at 37°C in a humid atmosphere with 5% CO2. After 4 h the number of viable cells in the 3D cell culture was recorded as time point 0 (T0), CellTiter-Glo® 3D Cell Viability Assay (Promega, cat. no. G9682) was added to the wells, following the manufacturer’s instructions and the contents transferred into a Corning® 96-well Flat Clear Bottom White (Corning, cat. no. 3610) for the reading with the Tecan Infinite® 200 Pro. After one week the measurement was repeated.

BE874 organoids were generated and expanded using a special lung cancer organoid (LCO) medium (File S5, Table 12).

BE874 organoids were transfected with ASOs as described for the NCI-H441. 24 h after transfection the cells were detached, counted and seeded onto Corning® 96-well Flat Clear Bottom White (Corning, cat. no. 3610) in 20 μl domes of LCO growth medium and Matrigel (1:1) with a density of 20,000 cells per dome. The Matrigel-containing PDX-organoids was allowed to solidify for an hour in the incubator at 37 °C before adding 80 μl LCO growth media on top. The organoids were allowed to grow in the incubator at 37°C in a humid atmosphere with 5% CO2. After 24h, 100 μl of CellTiter-Glo® 3D Cell Viability Assay (Promega cat. no. G9682) were added to the wells intended for the T0 and the luminescence was recorded with Tecan Infinite® 200 Pro. After three days the 80 μl of LCO media were added to the wells to keep them from drying out. After one week, the media was aspirated and replenished with fresh 80 μl, before proceeding with the measurement with CellTiter-Glo® 3D.

### Apoptosis assay

Annexin V and viability dye were used to detect early apoptotic and dead cells, respectively. 24 h after the transfection, cells were counted and 150,000 cells were re-suspend in 100 μl of 1X PBS. The viability dye (ThermoFisher, cat. no. 35111) was added (1:5,000) in 100 μl of 1X PBS and cells incubated for 30 minutes at 4°C. Cells were then washed once with 1X PBS and re-suspend in 100 μl of Annexin buffer PH 7, added PE Annexin (1:200; ThermoFisher, cat. no. L34960) and incubated 30 minutes at 4°C. After a wash with 1X PBS, cells were resuspended in 300 μl of Annexin buffer and underwent the flow analysis by using the LSR Fortessa instrument (BD Biosciences). Unstained cells were used as control.

### Cell cycle assay

Cells were transfected with ASOs using Lipofectamine3000 according to the manufacturer’s instructions. 24 h after, cells were harvested and fixed with 100 μl of BD Cytofix/Cytoperm Fixation (BD Biosciences, cat. no. 51-2090KZ) for 30 minutes at room temperature. The cells were then washed with 200 μl of 1X BD Perm/Wash (BD Biosciences, cat. no. 51-2091KE) and resuspended in 100 μl of 1X PBS. The K-i67 Antibody (ThermoFisher cat. no. 12-5698-82) was added (1:100) and incubated for 30 minutes at 4°C. Wash again with 1X BD perm/wash and stain with DAPI (Roche, cat. no. 10236276001) was added (1:10,000) in 100 μl of 1X PBS. Incubate 5’ at room temperature and wash with 1X PBS. Acquire the data with the Fortessa flow cytometer. Data analysis performed using FlowJo, and the different cell cycle phases were determined according to the Dean-Jett Fox (DJF) model.

### Low-throughput migration assay

Migration assay was performed as previously described(111). 24 h after ASOs transfection, A549 cells were counted and seeded in the upper part of Boyden chambers with the density of 35,000 cells/transwell. The upper part of transwell inserts was filled with media without FBS, while the lower part with media supplemented with 10% FBS to induce the directional movement of cells. After 24 h the cells were washed three times with 1X PBS and stained using 300 μl of crystal violet 1% for 30 minutes. Three washes with 1X PBS followed. The cells in the upper part of the membrane were removed by using a cotton swab. The chambers are left to dry overnight. The day after, the crystal violet was solubilized in 1X PBS containing 1% SDS and the absorbance at 595 nm was recorded by using the Tecan Infinite® 200 Pro.

### RNA isolation and qRT–PCR

To purify total RNAs from cultured cells, a Quick-RNA™ kit from ZymoResearch was used according to the manufacturer’s protocol. RNAs were reverse transcribed to produce cDNAs by using the GoScript™ Reverse Transcription System Kit (Promega, cat. no. A5003). The cDNAs were then used for qPCR to evaluate gene expression, using the GoTaq® qPCR Master Mix kit (Promega, cat. no. A6002). The expression of HPRT1 was used as an internal control for normalization. All the primers are listed in File S5, Table 13.

### RNA-sequencing and analysis

24 h after ASO transfection, A549 and H460 cells were harvested and the total RNA was extracted as explained before and samples’ quality was checked at Bioanalyzer. Libraries were prepared using the NEBNext® Ultra RNA Library Prep Kit and sequenced in paired-end 150 format to a depth of 30M reads/sample.

Transcript quantification was performed using Kallisto v0.46.0(112) against GENCODE v36(29). Gene level expression was inferred by aggregating the counts of the individual isoforms. Differential expression analysis was performed using Sleuth v0.30(113). Genes with a q-value <0.2 were considered significant. For CHiLL1, the genes that were significantly up- and down-regulated with two different ASO were selected. For CHiLL2, we selected the pool of common genes deregulated in A549 and H460.

Visualization of the results was produced in R 4.0.0 (R: The R Project for Statistical Computing, n.d.) using ggplot2 package v3.3.2 (Ggplot2 - Elegant Graphics for Data Analysis | Hadley Wickham | Springer, n.d.). Functional enrichment analysis was performed through the enrichR package v2.1(114).

### Fluorescent In-Situ Hybridization (FISH) and cell fractionation

FISH was performed on A549 cell lines, according to the Stellaris protocol (https://www.biosearchtech.com/support/resources/stellaris-protocols). For detection of CHiLL 1&2 at a single-cell level, pools of 25 and 48 FISH probes respectively were designed using the Stellaris probe designer software (www.biosearchtech.com). Cells were grown on round coverslip slides (ThermoFisher, 18mm), fixed in 3.7% formaldehyde and permeabilized in ethanol 70% overnight. Hybridization was carried out overnight at 37 °C in hybridization buffer from Stellaris. Cells were counterstained with DAPI and visualized using the DeltaVision microscope.

Nuclear and cytoplasmic fractionation was carried out in A549 cells as described previously(71). The pipeline used to analyse the data was adapted from CellProfiler (115), and it is named ‘SpeckleCounting’.

### ezTracks visualisation of RNA elements

Genomic tracks were retrieved for the hg38 human genome assembly from their original publications: predicted neo-functionalised fragments of transposable elements, also known as RIDLs (repeat insertion domains of lncRNA) (71), RNA structures conserved in vertebrates (CRS) (116), and ENCODE candidate cis-regulatory elements(117). In addition, the following tracks were downloaded from the UCSC Genome Browser(118): repeat-masked CpG islands, phastCons conserved elements in 7, 20, 30 and 100-way multiple alignments(119), and repeat families from the RepeatMasker annotation (Smit, AFA, Hubley, R & Green, P. RepeatMasker Open-4.0. 2013-2015; http://www.repeatmasker.org).

The comprehensive gene annotations for CHiLL1 (ENSG00000253616) and CHiLL2 (ENSG00000272808 and ENSG00000232386) loci were extracted from the GENCODE v36 GTF file(29). Then, all the exons corresponding to each locus were collapsed into a meta-transcript and output as separate GTF files using BEDTools merge(120). Third, configuration files for each locus were prepared to draw the meta-transcript annotation alongside the genomic tracks using the program ezTracks(82).

### Copy number analysis of pan-hallmark CRISPR candidates

The copy number status of A549 and in H460 cells was retrieved from the CCLE (https://portals.broadinstitute.org/ccle/data). Then, we intersected the hg19 coordinates of the pgRNA with the hg19 coordinates of the CCLE copy number data.

TCGA-LUAD copy number data were downloaded as ‘log2 ratio segment means’ using the R package TCGAbiolinks and converted to hg19 coordinates using liftOver. The values of each candidate were averaged across all TCGA-LUAD samples.

The pan-cancer recurrently amplified or deleted genomic regions were downloaded from the ICGC Data Portal (https://dcc.icgc.org/releases/PCAWG/consensus_cnv/GISTIC_analysis/all_lesions.conf_95.rmcnv.pt_170207.txt.gz). Then, we searched for overlaps between each candidate and the recurrent copy number altered regions (“Wide Peak Limits”). The differences in the proportions of amplified, deleted, or non-copy number altered hits versus non-hits were tested using Fisher’s exact tests.

## Supplementary information titles and legends

**Supplementary File 1.** Composition of CRISPR-del ‘‘libDECKO-NSCLC1’’ library. This file contains, per each target, the sequence of the pgRNAs and the target gene.

**Supplementary File 2.** Single screen results of all 10 screens using the CASPR pipeline.

**Supplementary File 3.** Results from the target prioritisation pipeline (TPP) (integration of all 10 screens (pan-hallmark), or split by phenotype (pro=proliferation; cis=cisplatin; mig=migration).

**Supplementary File 4.** List of plasmids used in this project. Related to the methods

**Supplementary File 5.** Tables 1-13. Sequence of oligonucleotides for, PCR qPCR, ASO, media composition and PCR condition.

## Supplementary figures

**Supplementary Fig. 1.** libDECKO-NSCLC1 library creation.

**Supplementary Fig. 2.** Assessing screen accuracy.

**Supplementary Fig. 3.** TPP quality assessment and CNV analysis. R

**Supplementary Fig. 4.** Tier 2 candidates and cancer hallmarks. R

**Supplementary Fig. 5.** Further information on CHiLL1 and CHiLL2.

**Supplementary Fig. 6.** CHiLL1&2 perturbation impacts disease transcriptome.

