## Supplementary figures and legends for "Multi-hallmark long noncoding RNA maps reveal non-small cell lung cancer vulnerabilities"

### Supplementary Fig. 1

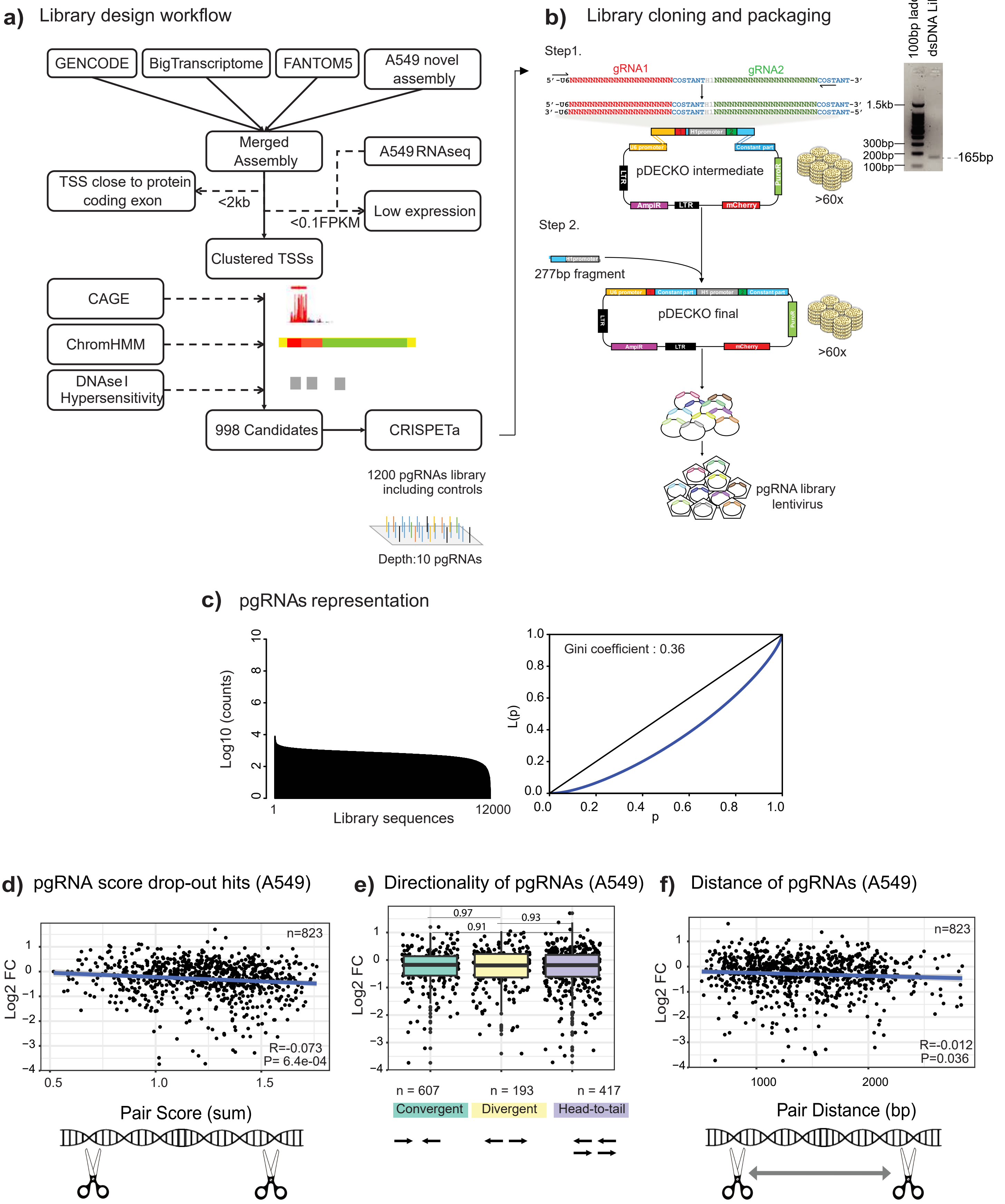

### Supplementary Fig. 2

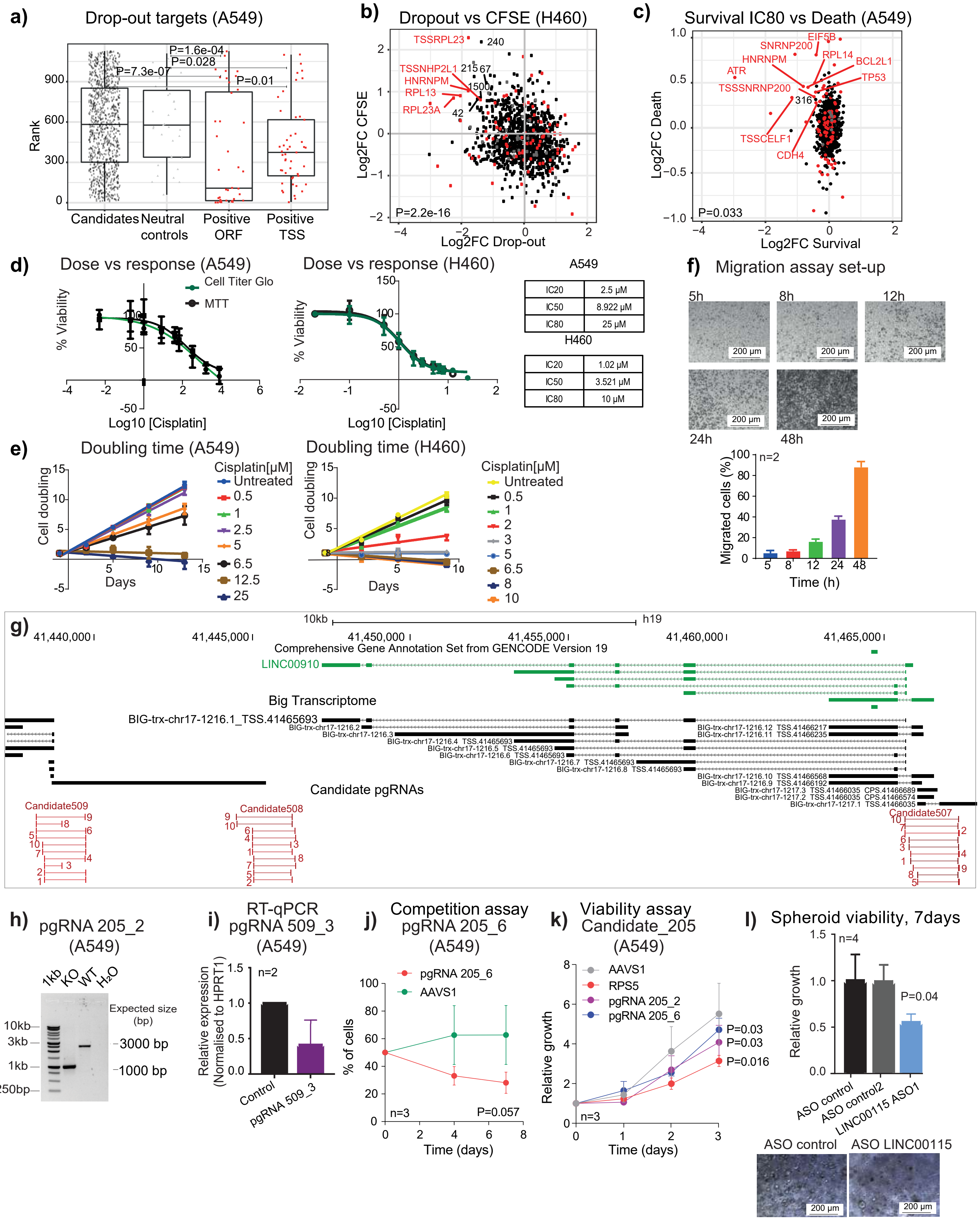

### Supplementary Fig. 3

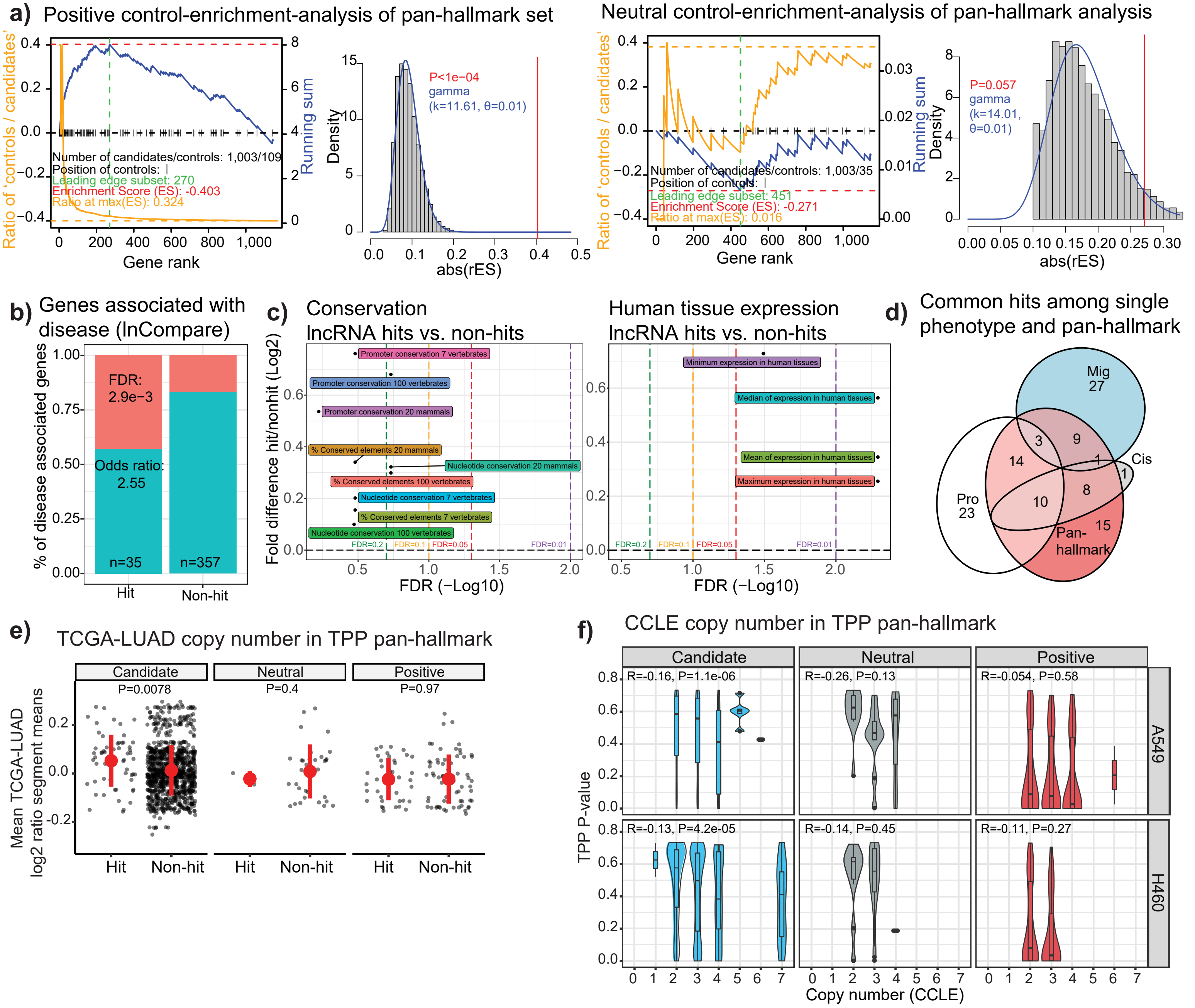

#### Supplementary Fig. 4

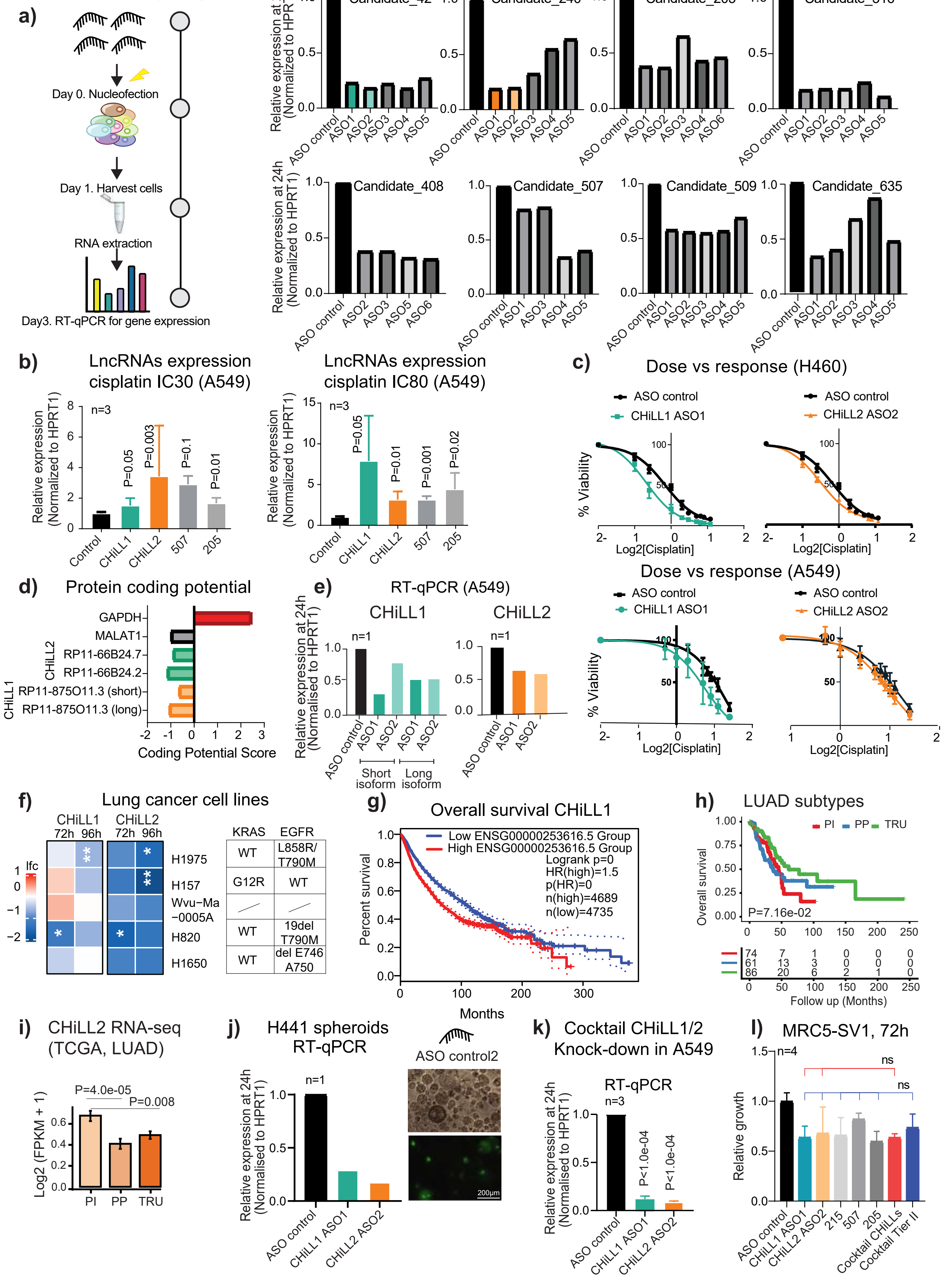

### Supplementary Fig. 5

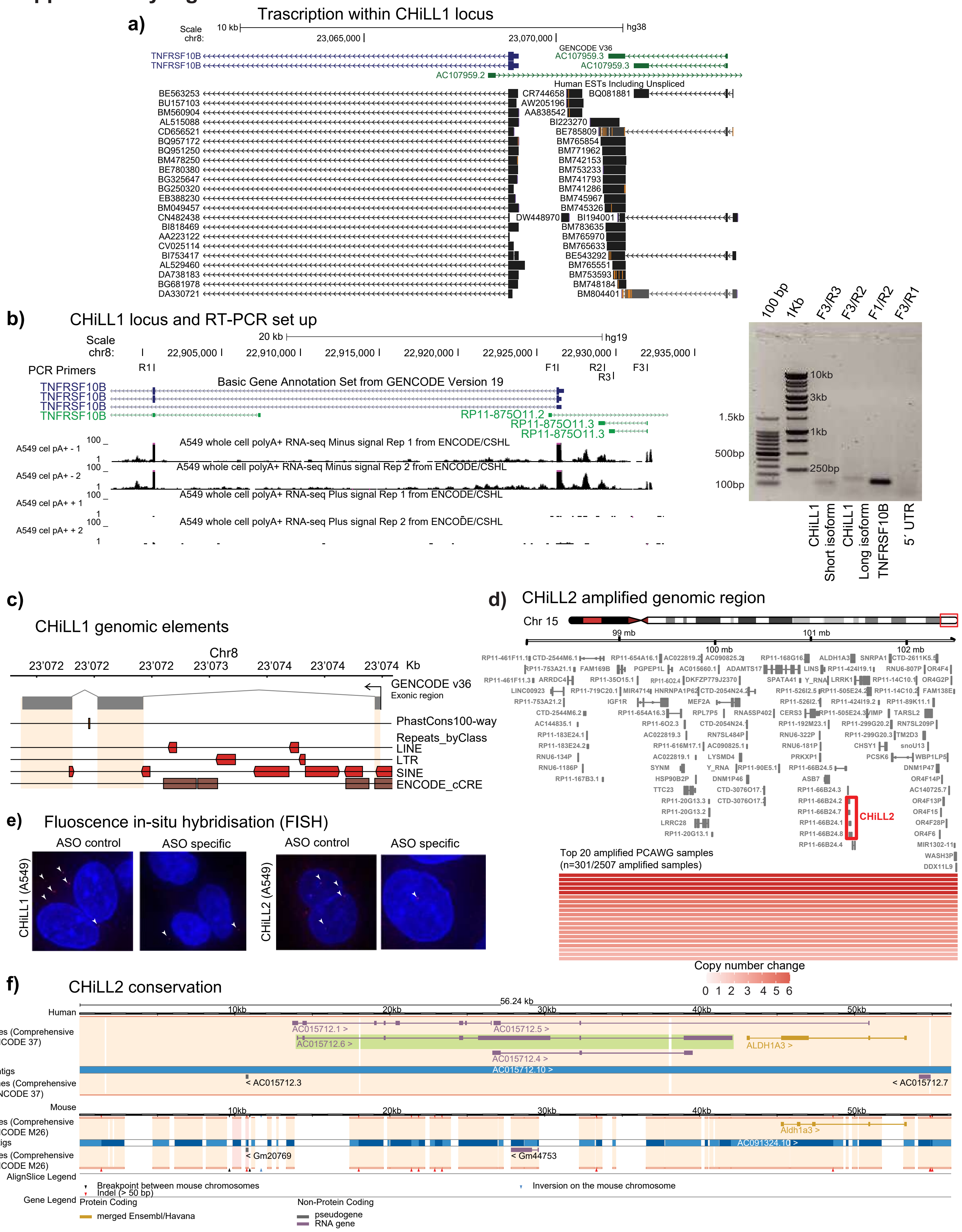

#### Supplementary Fig. 6

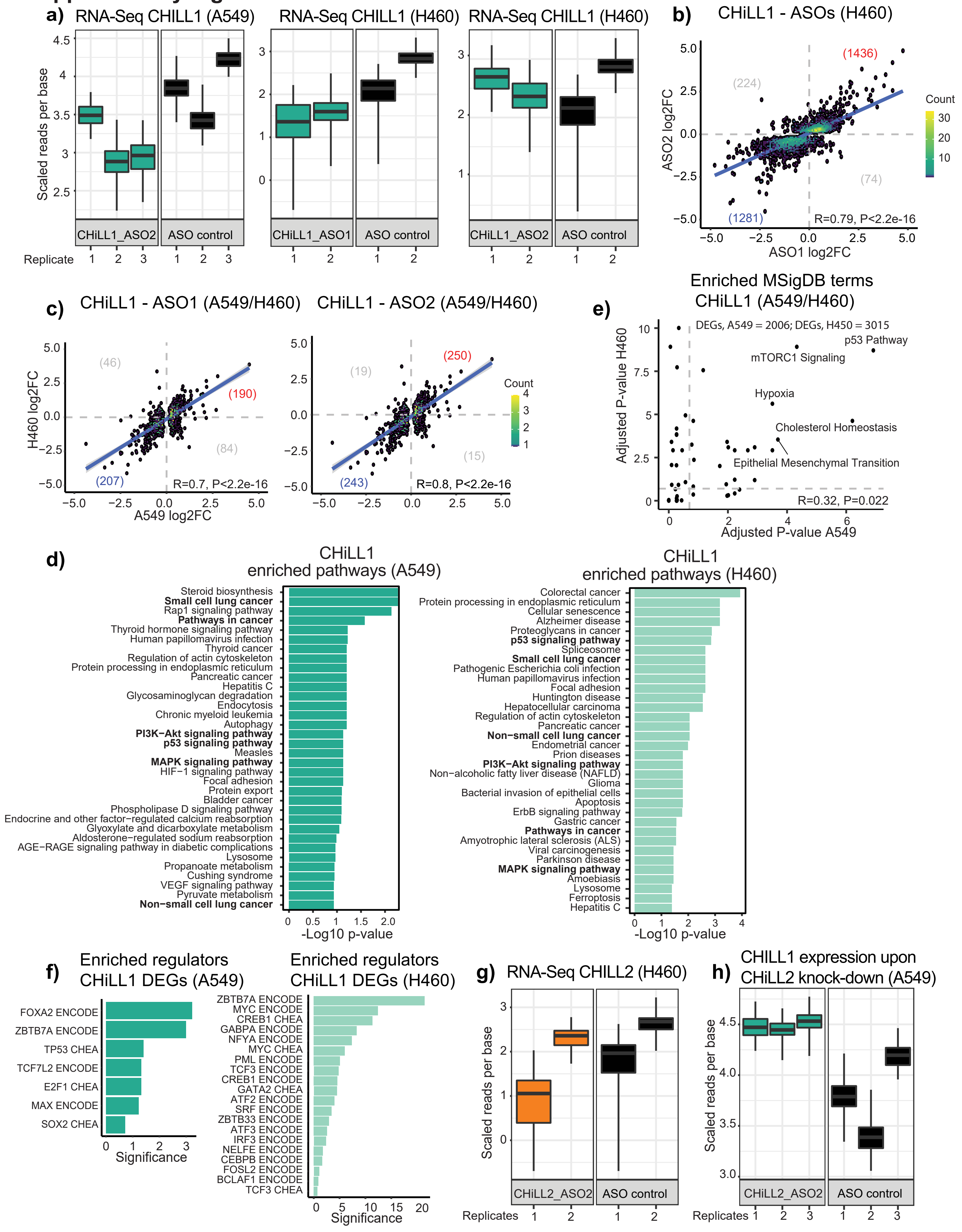

**Supplementary Fig. 1. libDECKO-NSCLC1 library creation.** **a**) Library design pipeline. LncRNAs from indicated annotations are merged and filtered by TSS proximity to protein-coding genes (<2 kb excluded) and by expression in A549 (<0.1 FPKM excluded). TSS are clustered together if closer than 300bp, then selected based on three evidence sources: CAGE, ChromHMM and DNAseI hypersensitivity. TSS candidates are targeted by 10 paired guide RNA (pgRNA) designs using CRISPETa. **b**) Library cloning. Oligonucleotide library of pgRNAs is amplified by PCR. Two steps of cloning follow to insert the amplified fragment and the constant part into the pDECKO backbone. Coverage of clones to library sequences was estimated to be >60x. The plasmid library is packaged into viral particles. **c**) Library representation. Left: y-axis: Number of reads per pgRNA sequence; x-axis: ranked pgRNAs from library. Less than 0.5% of the total pgRNAs constructs dropped out with no reads. The ratio of 10th – 90th percentile is 4.6-fold. Right: Lorenz curve, depicting library read coverage. Equality would be represented by the diagonal. **d**) Predicted deletion sizes of pgRNAs. Red line: mean; black line: median; candidates mean= 1528.432 bp; median=1512bp; controls: mean=1529.518b; median=1664.5bp. **e**) Figure depicts pgRNAs targeting lncRNA candidates defined as hits (A549 dropout, FDR<0.2). y-axis: log2 fold change in abundance; x-axis: sum of individual sgRNA scores for each pair from RuleSet2 algorithm. Significance estimated using linear model. **f**) As for (**e**), but separating pgRNAs according to the orientation of their individual sgRNAs. **g**) As for (**e**), but now for the genomic distance (bp) between the sgRNAs of each pgRNA.

**Supplementary Fig. 2. Assessing screen accuracy.** **a**) Each point represents a screen target, ranked by lowest P-value from A549 drop-out screen, so that position zero is most significant. Each positive control protein-coding gene is analysed separately using pgRNAs targeting the ORF and TSS. Statistical significance: Wilcoxon test. **b**) Correlation between negative drop-out and positive CFSE screens in H460 cells. Statistical significance: Pearson correlation. **c**) Correlation of negative survival (IC80) and positive death (IC20) screens in A549 cells. Statistical significance: Pearson correlation. **d**) Cisplatin dose-response curves in A549 and H460 cells. Error bars: standard deviation. **e**) Cell doublings calculated at indicated cisplatin concentrations. Error bars: standard deviation. **f**) Migration screen set-up. A549 cells (0.5M cells/well) were seeded onto transwell inserts and allowed to migrate for the indicated times. After crystal violet staining, cells that migrated through the membrane counted in five randomly selected fields. Results are expressed as means ± standard deviation (n=2). **g**) Genomic locus of Candidate_507, 508 and 509. **h**) Assaying genomic deletion with pgRNAs for Candidate_205: figure shows agarose gel electrophoresis of PCR product with primers amplifying the Candidate_205 target region. **i**) RT-qPCR with primers for Candidate_509 RNA. Error bars: standard deviation. **j**) Competition assay with an additional pgRNA targeting Candidate_205. The plot shows the percentage of mCherry+ or GFP+ cells at indicated times (n=3 biological replicates; error bar: standard deviation; two-tailed Student's *t* test). **k**) Cell viability assay with the two pgRNAs targeting candidate_205 (n=3 biological replicates; error bar: standard deviation; two-tailed Student's *t* test). The cell viability was measured by using the CellTiter-Glo® 2D. **l**) Cell viability in 3D spheroids grown from H441 cells upon ASO knockdown of LINC00115. The viability was measured with CellTiter-Glo® 3D, seven days after ASO transfection (n=4 biological replicates; error bars: standard deviation; statistical significance: one-tailed Student's *t* test).

**Supplementary Fig. 3. TPP quality assessment and CNV analysis. a**) Left: Enrichment of positive controls in the top of the TPP ranking results by gene set enrichment analysis (GSEA). The running sum (blue) is calculated by iterating through the ranks, increasing with positive controls, decreasing otherwise. The maximum absolute value of the running sum is the Enrichment Score (ES). Black ticks on the y-axis represent the location of positive controls. The significance of the enrichment is nominally evaluated by simulating 10,000 perturbations of the labelling of the genes. Right: The same analysis with neutral controls. **b**) Association of lncRNA pan-cancer hits and non-hits with disease lncRNAs as defined by the lnCompare database (hypergeometric test) [1]. **c**) The enrichment of evolutionary conservation (left) and healthy tissue expression (right). (Human Body Map project [www.illumina.com; ArrayExpress ID: E-MTAB-513]). Analysis was performed using lnCompare dataset [1]. Statistical significance: two-sided Wilcoxon test. **d**) The intersection of hits identified by TPP in the indicated datasets. **e**) Copy number status in TCGA-LUAD samples for pan-hallmark lncRNA hits. For each candidate or control, the log2 (ratio segment means) of each TCGA-LUAD sample was retrieved for the library target region. Then, the log2 ratio segment means across TCGA-LUAD were averaged for each candidate or control. Red dots and lines display the mean and the standard deviation, respectively. Statistical significance was estimated using Student's *t* tests. **f**)

Relationship between pan-hallmark TPP scores (uncorrected P-values) (y-axis) and copy number status (x-axis) in A549 or H460. Copy number status of the library target regions was retrieved from the Cancer Cell Line Encyclopedia (CCLE). Statistical significance: Pearson correlation.

**Supplementary Fig. 4. Tier 2 candidates and cancer hallmarks.** **a**) Left: Experimental strategy to test ASO knockdown effectiveness. Right: Measured gene knockdown by RT-qPCR in response to ASOs. **b**) RT-qPCR of Tier 2 candidates in A549 cells treated with cisplatin IC30 (left panel) and IC80 (right panel) for 72 h (n=3, error bars: standard deviation; statistical significance: one-tailed Student's *t* test). **c**) Dose response curves in H460 (upper panels) and A549 (lower panel) in cells transfected with ASO control and the best performing ASOs for CHiLL1 (left panels) and CHiLL2 (right panels). Error bars: standard deviation. **d**) Coding Potential Scores (CPS) of CHiLL 1&2 transcripts using the algorithm Coding Potential Calculator [2]. GAPDH and MALAT1 transcripts were used as reference for the protein-coding and non-protein coding, respectively. **e**) CHiLL1&2 knockdown efficiency measured by RT-qPCR, using two independent ASOs. **f**) The effect of CHiLL1&2 ASOs in additional NSCLC cell lines. Mutational status is indicated on the right. Columns: timepoints for the cell viability measurements. Values reflect the mean log2 fold change in viability following ASO transfection, with respect to a control non-targeting ASO, from at least two independent biological replicates. One-tailed Student's *t* test. * indicate P< 0.05; ** indicate P<0.01. **g**) Kaplan–Meier survival analysis of CHiLL1 including all the tumours available in the TCGA dataset. Analysis was performed using GEPIA2 tool with default settings [3]. **h**) Kaplan–Meier survival analysis of CHiLL2 in TCGA data within transcriptional subtypes. Proximal-inflammatory (PI), proximal-proliferative (PP) and terminal respiratory unit (TRU). **i**) CHiLL2 expression in transcriptional sub-types in TCGA dataset (n=513, Wilcoxon signed-rank tests). Proximal-inflammatory (PI), proximal-proliferative (PP) and terminal respiratory unit (TRU). **j**) Knockdown efficiency measured by RT-qPCR in 3D spheroids with CHiLL 1&2 ASOs relative to ASO control. Error bars: standard deviation. **k**) Knockdown efficiency measured in the cocktail experiment in A549 cells with CHiLL 1&2 ASOs relative to ASO control (n=3 biological replicates; error bars: standard deviation; statistical significance: two-tailed Student's *t* test). **l**) Growth of MRC5-SV1 cells in response to indicate ASO transfections (n=4 biological replicates; error bars: standard deviation; statistical significance: one-tailed Student's *t* test).

**Supplementary Fig. 5. Further information on CHiLL1 and CHiLL2.** **a**) CHiLL1 genomic locus. Note the support for CHiLL1 gene structure, and the lack of evidence for read through transcription between CHiLL1 and *TNFRSF10B,* from expressed sequence tags (ESTs). **b**) Searching for evidence of read through transcription between CHiLL1 and *TNFRSF10B.* Left: Expression of RNA from CHiLL1 locus, and primers used. Right: Agarose gel electrophoresis of RT-PCR products using indicated primers with A549 cDNA. **c**) Genomic elements in the CHiLL1 locus. GENCODE v36 exons were merged into a single transcript and queried against a set of genome-wide element annotations using ezTracks [4]. **d**) Recurrently amplified genomic region that encompasses CHiLL2 according to Pan-Cancer Analysis of Whole Genomes (PCAWG) Consortium. Ensembl 75 (hg19) annotation was used. For each gene, the longest transcript is shown. CHiLL2 locus is highlighted. The copy number gain for the top 20 amplified samples in PCAWG is shown. **e**) Representative confocal microscopy images of RNA-FISH performed with CHiLL1, CHiLL2 probe sets upon treatment with targeting ASOs or non-targeting ASO control in A549 cells. Selected lncRNA foci in the treated sample and in the control are arrowed. **f**) CHiLL2 orthology between human annotation (GENCODE 37) and mouse (GENCODE M26).

**Supplementary Fig. 6. CHiLL1&2 perturbation impacts disease transcriptome. a**) Response of CHiLL1 expression to ASO transfection, as measured by RNA-seq. The y-axis represents the normalized expression (counts) per nucleotide and the boxplots show the variance of inference using bootstraps from Kallisto. **b**) Changes in gene expression (log2 fold change) with two different CHiLL1 ASOs in H460. Numbers indicate the differentially expressed genes in each part of the scatter plot. Statistical significance: Pearson correlation. **c**) As for (b), but comparing effects of the same CHiLL1 ASO in two cell backgrounds (left: ASO1; right; ASO2). **d**) KEGG pathways enriched for CHiLL1 target genes, for A549 (left) and H460 (right) cells. Analysis was performed using common differentially expressed genes between the ASO1 and ASO2 knockdown. **e**) Gene ontology analysis performed using the Molecular Signatures Database (MSigDB). Shown are enriched terms between A549 and H460 cells by using the set of common differentially-expressed genes of ASO1&2 for CHiLL1.

Statistical significance: Pearson correlation. **f**) Binding sites [5,6] enriched in the intersection of differentially expressed genes (DEGs) from ASO1&2 knockdown of CHiLL1. **g**) Expression of CHiLL2 in H460 in response to CHiLL2 ASOs. The y-axis represents the normalized expression (counts) per nucleotide and the boxplots show the variance of inference using the bootstraps of kallisto. **h**) Expression of CHiLL1 (RNA-seq) upon ASO knockdown of CHiLL2. The y-axis represents the normalized expression (counts) per nucleotide and the boxplots show the variance of inference using the bootstraps of kallisto.
