## Supplementary.File5 for "Multi-hallmark long noncoding RNA maps reveal non-small cell lung cancer vulnerabilities"

| **Table 1. Validated candidates and ASOs** (- not annotated in Gen v27) | | | | n. |
|---|---|---|---|---|
| **Candidate** | **Name** | **Annotation** | **ASO Sequences** | |
| 42 | RP11-875O11.3 | ENSG00000253616 | CAGGAGAAAAGCACAC | 3 |
| | | | ATTCTGGGTCACTGCT | 5 |
| 240 | RP11-66B24 | ENSG00000272808 | CATAATCTGGGAACGA | 1 |
| | | | GTGTGGTTGGAAGCTA | 3 |
| 205B | LINC00115 | ENSG00000225880 | CAGAAGCACGAGGGTT | 1 |
| | | | AAGCTGAACCTGACAC | 2 |
| 205D | LINC01128 | ENSG00000228794.10 | ACATAGATTCATCGTG | 1 |
| 205D | LINC01128 | ENSG00000228794.10 | CTGAAGACCGCAGTTA | 2 |
| 507 | LINC00910 | ENSG00000188825.9 | CCTTTGCGGACAGTTG | 1 |
| | | | CTGATGACAGGAGTTA | 5 |
| 215 | AC104024.3 | ENSG00000287114 | TCGTCCAGCTAATAAT | 1 |
| | | | TTGGACAGAGTAAGCA | 4 |
| 489 | MIR23AHG (v.38) not annotated in v.19 | ENSG00000267519.6 (v.38) | CCAGGGACGAGAACGG | 1 |
| | | | CGCATTGGGAGACTAA | 3 |
| 509 | | - | GGTTGAATCGATTGGA | 2 |
| | | | CGTAAAGGTGCAAACG | 3 |
| 316 | RP11-222A11.1 | ENSG00000228065.6 | GACATGCAGAGAGTTA | 1 |
| | | | AAAACGAGGTCTAGTA | 2 |
| 408 | | - | CATGAGAGAAGAATCT | 4 |
| | | | CAATTTACCCTCAATC | 5 |
| 311 | RP11-137H2.4 | ENSG00000226659 | TAAACAATTAGGCGTG | 1 |
| | | | AGATGCGAGAGTGACT | 4 |
| 635 | HOXA-AS3 | ENSG00000254369 | CACGAGTGAAGAGCAT | 1 |
| | | | GCGTGTGCACTAACAT | 2 |
| 448 | CECR7 | ENSG00000237438.2 | | |
| MALAT1 | | | Qiagen control | |
| MTOR | | | Qiagen control | |
| Control A | | | Qiagen neutral control | |

| **Table 2. CRISPETA runs** | | | | | | | | |
|---|---|---|---|---|---|---|---|---|
| | Run1 | Run2 | Run3 | Run4 | Run5 | Run6 | Run7 | Tun8 |
| -eu | 0 | | | | | | | |
| -ed | 0 | | | | | | | |
| -du | 1000 | | 1200 | | | | | |
| -dd | 1000 | | 1200 | | | | | |
| -si | 0.2 | | | | | | | 0 |
| -t | 0,0,0,x,x | 0,0,1,x,x | | 0,0,5,x,x | 0,1,5,x,x | 0,x,x,x,x | 1,x,x,x,x | 1,x,x,x,x |

| **Table 3. PCR reaction ssDNA to dsDNA** | |
|---|---|
| Component | Amount per reaction |
| 5x Buffer | 20 ul |
| dNTPs 10 mM | 3 ul |
| Forward oligo 100uM | 2 ul |
| Reverse oligo 100 uM | 2 ul |
| Template DNA 10ng/ul | 2 ul |
| Phusion Polymerase High Fidelity 2U/ul | 2ul |
| UltraPure water | Up to 100 ul |

| **Table 4. pDECKO_backbone plasmid digestion** | |
|---|---|
| Component | Amount per reaction |
| pDECKO plasmid | 5 ug |
| Tango 10x Buffer | 5 ul |
| DTT (20mM) | 2.5 ul |
| Esp3I (BsmBI) | 1 ul |
| UltraPure H2O | Up to 50 ul |

| **Table 5. Long oligos to generate the constant part (HPLC purification)** | |
|---|---|
| Oligos | Sequences (5'-3') |
| Forward_1 (F1) 125 bp | TAGAAATAGCAAGTTAAAATAAGGCTAGTCCGTTATCAACTTGAAAAAGTGGCACCGAGTCGGTGCTTTTTTGAACGCTGACGTCATCAACCCGCTCCAAGGAATCGCGGGCCCAGTGTCACTAG |
| Forward_2 (F2) 148 bp | GCGGGAACACCCAGCGCGCGTGCGCCCTGGCAGGAAGATGGCTGTGAGGGACAGGGGAGTGGCGCCCTGCAATATTTGCATGTCGCTATGTGTTCTGGGAAATCACCATAAACGTGAAATGTCTTTGGATTTGGGAGTCTTATAAGTT |
| Reverse_1 (R1) | GCGCACGCGCGCTGGGTGTTCCCGCCTAGTGACACTGGGCCCGCGATTCCTTGGAGCGGGTTGATGACGTCAGCGTTCAAAAAAGCACCGACTCG |

| | |
|---|---|
| 146 bp | GTGCCACTTTTTCAAGTTGATAACGGACTAGCCTTATTTTAACTTGCTAT T |
| Reverse_2 (R2) 127 bp | ACAGAACTTATAAGACTCCCAAATCCAAAGACATTTCACGTTTATGGTGA TTTCCCAGAACACATAGCGACATGCAAATATTGCAGGGCGCCACTCCC CTGTCCCTCACAGCCATCTTCCTGCCAGG |

| **Table 6. Colony PCR primers** | |
|---|---|
| Primer | Sequences (5'-3') |
| Oligo_colony_Fw | GTACAAAATACGTGACGTAG |
| Oligo_colony_Rv | ATGTCTACTATTCTTTCCCC |

| **Table 7. Colony PCR reaction**<br>95ºC 2', 95ºC 30'', 60ºC 40'', 72ºC 2' for 29 cycles, 72ºC 2', 4ºC forever | |
|---|---|
| Component | Amount per reaction |
| 5x Buffer | 10 ul |
| dNTPs 10 mM | 1 ul |
| Oligo_colony_Fw 10 uM | 1 ul |
| Oligo_colony_Rv 10 uM | 1 ul |
| Template colony | 2 ul |
| Go Taq G2 polimerase (Promega) | 0.25 ul |
| Nuclease-free water | Up to 50 ul |

| **Table 8. Primers for NGS sequencing of the library (HPLC purification)** | |
|---|---|
| Primer | Sequences (5'-3') |
| F (staggered oligo 0) | TTCAGACGTGTGCTCTTCCGATCTGTGGAAAGGACGAAACACCg |
| R (staggered oligo 0) | CCCTACACGACGCTCTTCCGATCTcaagatctagttacgccaagcttAAA |
| S1_F (Staggered ) | TTCAGACGTGTGCTCTTCCGATCTHGTGGAAAGGACGAAACACCg |
| S2_F (Staggered ) | TTCAGACGTGTGCTCTTCCGATCTHMGTGGAAAGGACGAAACACCg |
| S3_F (Staggered ) | TTCAGACGTGTGCTCTTCCGATCTHMMGTGGAAAGGACGAAACACCg |

| | |
|---|---|
| S4_F (Staggered) | TTCAGACGTGTGCTCTTCCGATCTNNMMGTGGAAAGGACGAAACACCg |
| S5_F (Staggered) | TTCAGACGTGTGCTCTTCCGATCTNNMMCGTGGAAAGGACGAAACACCg |
| S1_R (Staggered) | CCCTACACGACGCTCTTCCGATCTDcaagatctagttacgccaagcttAAA |
| S2_R (Staggered) | CCCTACACGACGCTCTTCCGATCTDKcaagatctagttacgccaagcttAAA |
| S3_R (Staggered) | CCCTACACGACGCTCTTCCGATCTDKKcaagatctagttacgccaagcttAAA |
| S4_R (Staggered) | CCCTACACGACGCTCTTCCGATCTNNKTcaagatctagttacgccaagcttAAA |
| S5_R (Staggered) | CCCTACACGACGCTCTTCCGATCTNNBBTcaagatctagttacgccaagcttAAA |

**Table 9. Forward staggered oligos for PCR and NGS of the library** (HPLC purification)
The 6 different NGS_FwSt primers contain 1–6 additional nucleotides after the P5 Illumina adapter designed to increase the diversity of the NGS library.
Primer structure as follows: P5 – linker –SBS3 sequencing primer, read 1 – **staggered nucleotides** – *template primer for p-DECKO vector forward*

| Primer | Sequences (5’-3’) |
|---|---|
| NGS_FwSt_1 | AATGATACGGCGACCACCGAGATCTACACTCTTTCCCTACACGACGCTCTTCCGATCT*GTGGAAAGGACGAAACACCg* |
| NGS_FwSt_2 | AATGATACGGCGACCACCGAGATCTACACTCTTTCCCTACACGACGCTCTTCCGATCT**H***GTGGAAAGGACGAAACACCg* |
| NGS_FwSt_3 | AATGATACGGCGACCACCGAGATCTACACTCTTTCCCTACACGACGCTCTTCCGATCT**HM***GTGGAAAGGACGAAACACCg* |
| NGS_FwSt_4 | AATGATACGGCGACCACCGAGATCTACACTCTTTCCCTACACGACGCTCTTCCGATCT**HMM***GTGGAAAGGACGAAACACCg* |
| NGS_FwSt_5 | AATGATACGGCGACCACCGAGATCTACACTCTTTCCCTACACGACGCTCTTCCGATCT**NNMM***GTGGAAAGGACGAAACACCg* |
| NGS_FwSt_6 | AATGATACGGCGACCACCGAGATCTACACTCTTTCCCTACACGACGCTCTTCCGATCT**NNMMC***GTGGAAAGGACGAAACACCg* |

**Table 10. Reverse barcoded oligos for PCR and NGS of the library** (HPLC purification)
Primer structure as follows: P7 – linker – **barcode** – SBS12 sequencing primer, read 2 – *template primer for p-DECKO vector reverse.*

| example | Sequences (5’-3’) |
|---|---|
| NGS_RV_Barcode_1 | CAAGCAGAAGACGGCATACGAGAT**NNNNNNNN**GTGACTGGAGTTCAGACGTGTGCTCTTCCGATCT*CAAGATCTAGTTACGCCAAGCTTAAA* |

| **Table 11. Primers for the genomic deletion** | |
|---|---|
| gDNA_Candidate331_For | TTGCAACCCCCAAACAGACT |
| gDNA_Candidate331_Rev | GGGGCACCATTTTGGACCTA |
| gDNA_Candidate205_For | AGCCTGTCACAAACTGATTCTTA |
| gDNA_Candidate205_Rev | TTGTTGACCCGGAAACGGAT |

| **Table 12. Composition of lung cancer organoids medium** | | | |
|---|---|---|---|
| Substances | Final Conc. | Supplier | Catalogue number |
| DMEM:F-12 HAM medium | - | Sigma-Aldrich | D6421 |
| EGF | 50 ng/mL | Thermo Fisher Scientific | PHG6045 |
| FGF | 20 ng/mL | Thermo Fisher Scientific | PHG6015 |
| L-Glutamine | 2 mM | Sigma-Aldrich | G7513-100mL |
| OmniPur HEPES | 10 mM | Sigma-Aldrich | 5320-500GM |
| Penicillin/Streptomycin | 100 U/mL | Sigma-Aldrich | P3032-10MU S6501-25G |
| N-2 supplement | 1x | Thermo Fisher Scientific | 17502-048 |
| B-27 supplement | 1x | Thermo Fisher Scientific | 17504-044 |
| Noggin | 100 ng/mL | Prospec | CYT-475 |
| ROCK-inhibitor (Y-27632) | 10 µM | Stemcell | 72304 |

| Table 13. Primers RT-qPCR | |
|---|---|
| HPRT1_For | ATGACCAGTCAACAGGGGACAT |
| HPRT1_Rev | CAACACTTCGTGGGGTCCTTTTCA |
| Candidate_331_For | CAGGGAGCAGGGACTATCAA |
| Candidate_331_Rev | TGGTCTTCCAACATGGGCTTG |
| Candidate_205_For (LINC00115) | CCTAGTTCTCTTCACCGTCCG |
| Candidate_205_Rev (LINC00115) | AAGACAAGCCACATGCCGAA |
| Candidate_205_For (LINC01128) | AGAGGTTAAAAGTCACAAGGGTGT |
| Candidate_205_Rev (LINC01128) | GCCTTGACAGCAAGCCTAGA |
| Candidate_42_For (ENST00000520840.2) | GCAGTGACCCAGAATGAGGAAG |
| Candidate_42_Rev (ENST00000520840.2) | TACTGAAATTGGAGGCTGTGGA |
| Candidate_42_For (ENST00000523806.1) | GCAGTGACCCAGAATGAGGT |
| Candidate_42_Rev (ENST00000523806.1) | GCTCTAGCTTCCAGGTTGGG |
| Candidate_240_For | CTCACGGCAGCTATGAGACT |
| Candidate_240_Rev | GCTCCAAGATGCCACTCACA |
| Candidate_448_For | GAAACCTCCTCGACACTCCG |
| Candidate_448_Rev | AGTCTTCGAACAGGCTGCAA |
| Candidate_215_For | GCAATTGTACCTGAGGACCCA |
| Candidate_215_Rev | TGGCATATGGTGGATGTTCCC |
| Candidate_489_For | AAGCGCTCATTCAAGGTTGC |
| Candidate_489_Rev | GGTTCAGTCTGGGCCCTTTT |
| Candidate_408_For | GCGATGGAAGAAGTTTCGCC |
| Candidate_408_Rev | GGAACTCAGGTAACAGGAATTTCAC |
| Candidate_316_For | GACCAACTCCGTTTCCCGAT |
| Candidate_316_Rev | TCAAGGGCCCAGCCTTATTC |
| Candidate_635_For | AATTCCACCCACGCACCTAT |

| | |
|---|---|
| Candidate_635_Rev | GAGCCACCGTTAATTCAGCC |
| Candidate_507_For | TCCTTGCTAACCACACGGAC |
| Candidate_507_Rev | ATGTGGGTCCCAGTATCCGA |
| Candidate_509_For | TTGGCACACTCAGATGCGAT |
| Candidate_509_Rev | AAAACAGTCCCGCTTGGGAT |
| GAPDH_For | GCACCGTCAAGGCTGAGAAC |
| GAPDH_Rev | TGGTGAAGACGCCAGTGGA |
| MALAT1_For | GATTGAGGCGTTTTCCAAGA |
| MALAT1_Rev | ACTTTCTCCCCCAACTGCTT |
